# SMURF2/USP7-mediated ubiquitination of KAP1 controls its SUMO E3 ligase activity and chromatin regulation

**DOI:** 10.1101/2025.08.12.669848

**Authors:** Sandy Boutros-Suleiman, Kavya Nambiar, Pooja Anil Shah, Nilabja Roy Chowdhury, Meir Shamay, Gal Levy-Cohen, Michael Blank

## Abstract

KRAB-associated protein 1 (KAP1) is a critical nuclear protein that regulates chromatin architecture and gene expression, primarily through its SUMOylation activity. However, the mechanisms controlling KAP1 remain poorly understood. In a recent study, we identified the E3 ubiquitin ligase SMURF2 as a direct interactor and ubiquitin ligase for KAP1. Here, we show that SMURF2-mediated ubiquitination of KAP1 at lysines K254, K319, and K779 regulates its SUMOylation activity. We further demonstrate that SMURF2 collaborates with the deubiquitinase USP7/HAUSP, recruiting it to KAP1, to regulate a dynamic ubiquitination-deubiquitination cycle at these key residues. Cells expressing a KAP1 mutant, resistant to SMURF2-USP7 regulation, exhibit profound alterations in chromatin structure, gene expression, protein-protein interactions, and elevated LINE-1 retrotransposon activity. Notably, disrupting the SMURF2-USP7-KAP1-SUMOylation axis in cancer cells reduces their malignancy and tumor growth in vivo. Collectively, these findings uncover a novel regulatory mechanism for KAP1, highlighting its pivotal role in orchestrating critical cellular processes.

## INTRODUCTION

KAP1 (Krüppel-associated box (KRAB)-associated protein 1), also known as TRIM28 (tripartite motif-containing protein 28) and TIF1β (transcription intermediary factor 1β), is a ubiquitously expressed protein with versatile cellular functions. It is involved in the regulation of the epigenetic landscape, chromatin structure and topology, cellular and viral gene expression programs, including the expression of endogenous retroviruses (ERVs), as well as DNA damage response and maintenance of genomic integrity^1–5^. KAP1 regulates gene expression of both protein-coding and non-coding RNAs, such as long non-coding RNAs (lncRNAs), and LINE-1 (long interspersed nuclear elements) retrotransposons, which play critical roles in regulating chromatin structure and gene expression by influencing transcription, RNA splicing, stability, and translation^6–8^. By regulating these processes, KAP1 influences multiple aspects of cell physiology, including cell differentiation, renewal, survival, and death. It also plays a significant role in pathobiology, especially in cancer, where KAP1 is primarily considered a tumor-promoting factor, although its tumor-suppressive role has also been documented, particularly in Wilms’ tumors^9–13^.

KAP1 exerts its cellular functions through several distinct mechanisms: (i) acting as a SUMO E3 ligase, KAP1 SUMOylates itself^14,15^, enabling it to associate with the SUMO-interacting motifs found in many chromatin-modifying enzymes and associated proteins. These include the nucleosome remodeling and deacetylase (NuRD) complex^16^, the histone H3 lysine methyltransferase SETDB1^17^, and heterochromatin-associated proteins such as HP1^18–20^, which together establish a local repressive heterochromatin environment. The specificity of KAP1’s targeting to the genome for heterochromatin deposition and maintenance (e.g., in the repression of ERVs) is enabled through KAP1’s interactions with the KRAB-domain-containing zinc-finger DNA binding proteins (KRAB-ZFPs)^21,22^. (ii) KAP1-mediated SUMOylation of other proteins, such as IRF7^23^, Vps34^24^, PCNA^25^, CDK9^26^, NLRP3^27^, NPM1/B23^28^, and PD-L1^29^, affecting their functions. (iii) Acting as an ubiquitin E3 ligase (RING-type), KAP1 has been reported to ubiquitinate and regulate the stability of several cellular proteins, including BCL2A1^30^, cyclin A and TFIIB^31^, pRB^32^, as well as AMPK following the formation of a *de novo* complex between KAP1 and MAGE proteins^33^. (iv) Through extensive protein-protein interactions (PPIs), KAP1 is involved in the regulation of multiple cellular functions, with the most documented role in DNA repair and gene expression. Noteworthy, in gene expression regulation, KAP1 was reported to function both as a transcriptional repressor (co-repressor) and transcriptional activator, through distinct mechanisms^4^.

Despite the remarkable progress in elucidating the role of KAP1 in normal cell biology (both during development and in adulthood) and in several disease states, the regulatory mechanisms of KAP1 remain poorly understood.

Recently, we identified the E3 ubiquitin ligase SMURF2 (Smad Ubiquitin Regulatory Factor 2) as a direct interactor and ubiquitin ligase for KAP1, demonstrating an important role for SMURF2’s catalytic activity in regulating KAP1’s PPIs^34^. In the present study, we show that SMURF2-mediated ubiquitination of KAP1 at lysines K254, K319, and K779 regulates its SUMOylation activity. We further demonstrate that SMURF2 works in tandem with the deubiquitinase USP7, also known as HAUSP (Herpesvirus-Associated Ubiquitin-Specific Protease), recruiting it to KAP1 to regulate its ubiquitination-deubiquitination cycle at these lysine residues. Disruption of this regulatory axis leads to significant alterations in epigenetic landscape, chromatin structure, gene expression (including elevated expression of LINE-1 retrotransposons), and changes in the KAP1 interactome, affecting critical cellular processes both in vitro and in vivo.

## RESULTS

### Identification of SMURF2-mediated ubiquitination sites on KAP1

To identify the KAP1 residues undergoing SMURF2-dependent ubiquitination, we employed three complementary approaches, incorporating both in vitro (using purified proteins) and in-cellulo KAP1 ubiquitination assays, utilizing either wild-type SMURF2 (SMURF2WT) or its E3-ligase-dead mutant (SMURF2^C716G^; SMURF2*Mut*).

In approach #1 (Fig. 1A), an in vitro ubiquitination reconstitution assay was performed using purified components: the ubiquitin-activating enzyme (E1), the ubiquitin-conjugating enzyme (E2), HA-ubiquitin, and either wild-type or catalytically mutant GST-SMURF2, along with MYC-tagged KAP1. Purified GST protein was included as an additional control. After incubation, MYC-KAP1 was pulled down from the reaction, resolved by SDS-PAGE, and stained with Coomassie. The bands corresponding to KAP1 and its potential ubiquitination were excised from the gel and subjected to LC-MS/MS. SMURF2-mediated ubiquitination of KAP1 and its dependence on SMURF2’s catalytic activity were confirmed by immunoblots (Fig. 1B). In approach #2 (Fig. 1C), FLAG-KAP1 was expressed in HEK-293T cells for an in-cellulo ubiquitination assay and subsequently affinity-purified using an anti-FLAG antibody. The samples were then electrophoresed, stained with Coomassie, and the bands corresponding to KAP1 and its potential modifications were excised and submitted to LC-MS/MS, as in approach #1. Western blot analysis confirmed the in-cellulo ubiquitination of KAP1 by catalytically active SMURF2 (Fig. 1D, lane 4 vs. lanes 3 and 5).

**Figure 1.**
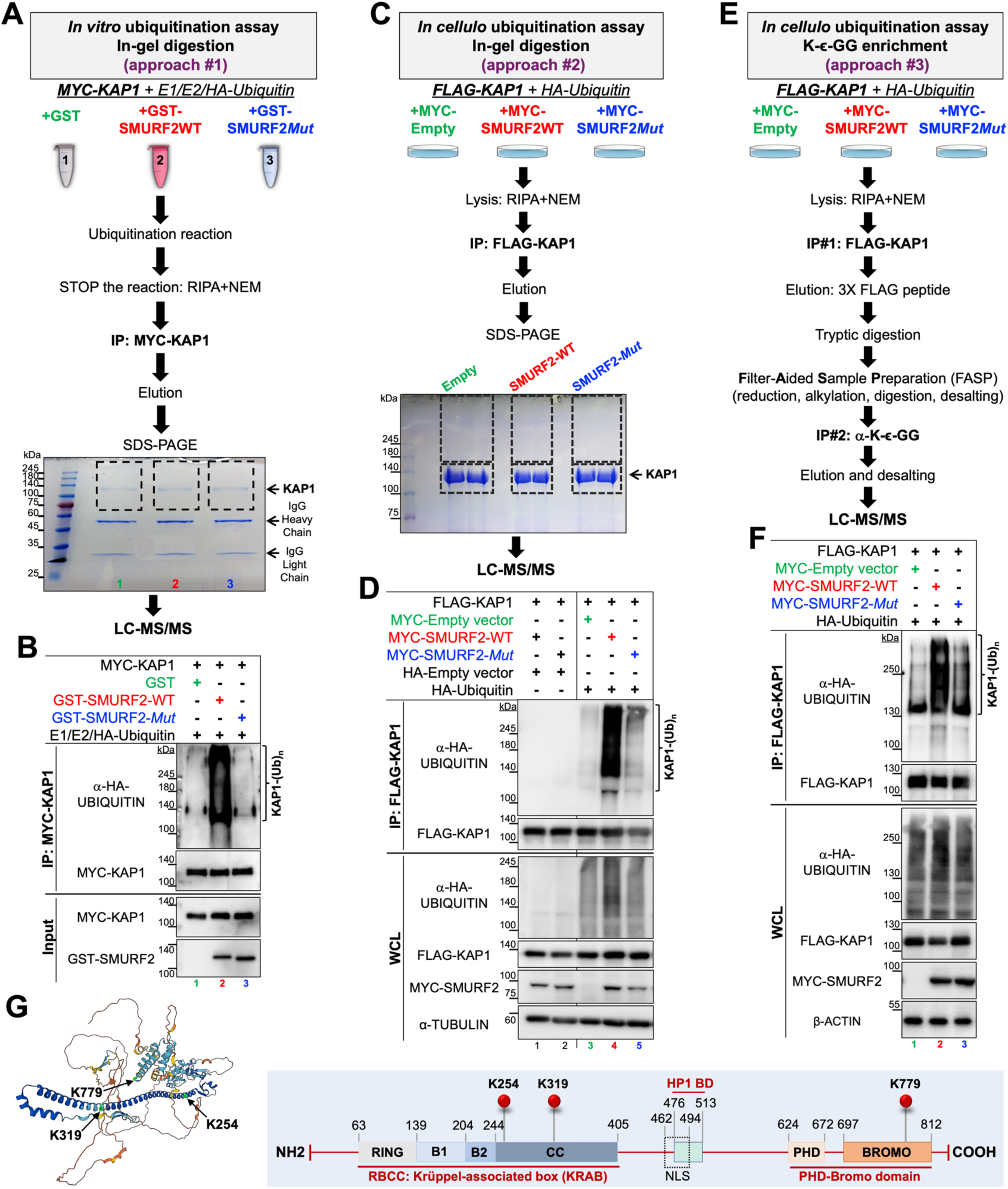
Identification of SMURF2-mediated ubiquitination sites on KAP1. **(A)** Flow chart of Approach #1 (in vitro ubiquitination assay) used to map the ubiquitination sites on KAP1. The dotted lines on the Coomassie-stained gel indicate the regions corresponding to KAP1 and its potential ubiquitination, which were excised and subjected to MS analysis. **(B)** Western blot analysis of the samples described in panel (A) showing ubiquitination of purified MYC-KAP1 by catalytically-active SMURF2 (GST-SMURF2-WT). Ubiquitination is absent when KAP1 is co-incubated with either the E3 ligase-dead SMURF2^C716G^ (GST-SMURF2-*Mut;* lane 3 vs. lane 2) or with GST (lane 1 vs. lane 2). **(C)** Workflow of Approach #2 (in-cellulo ubiquitination assay) performed in HEK-293T cells transiently co-expressing FLAG-KAP1, MYC-SMURF2 (either WT or *Mut*) and HA-Ubiquitin. The dotted lines on the Coomassie-stained gel indicate the regions excised for MS analysis, performed on duplicate samples. To minimize interference from unmodified KAP1, the samples were excised and submitted for LC-MS/MS separately, as indicated. **(D)** Western blot analysis of KAP1 ubiquitination by E3 ligase-proficient SMURF2 in HEK-293T cells, corresponding to the workflow in panel (C). FLAG-KAP1 was immunoprecipitated using anti-FLAG resin, and ubiquitination was detected by immunoblotting with anti-HA (ubiquitin) antibody. The specificity of the detection is demonstrated by the absence of ubiquitination signals in FLAG-KAP1 samples co-transfected with an HA-empty vector (lanes 1 and 2), compared to FLAG-KAP1 samples co-transfected with HA-Ubiquitin (lanes 3-5). The dependence of KAP1 ubiquitination on catalytic activity of SMURF2 versus its E3 ligase-dead form is also shown. IP: immunoprecipitation. WCL, whole cell lysates. **(E)** Flow chart of Approach #3: In-cellulo ubiquitination assay in HEK-293T cells combined with GG remnant (K-ε-GG) immunoaffinity purification to enrich for KAP1-ubiquitinated peptides. **(F)** Western blot analysis corresponding to the workflow in panel (E), validating SMURF2-mediated ubiquitination of KAP1 in the samples. **(G)** Identified lysine residues of KAP1 as ubiquitination sites of SMURF2. The left panel shows the mapping of these residues in the predicted full-length structure of human KAP1/TRIM28 (AlphaFold model AF-Q13263-F1; https://alphafold.ebi.ac.uk/entry/Q13263). Color coding indicates per-residue model confidence scores (pLDDT) as follows: dark blue, very high confidence (pLDDT > 90); light blue, high confidence (90 > pLDDT > 70); yellow, low confidence (70 > pLDDT > 50); orange, very low confidence (pLDDT < 50). The right panel shows a schematic diagram indicating the positions of the identified lysine residues within annotated KAP1 domains.

In approach #3 (Fig. 1E), FLAG-KAP1 was also affinity-captured from an in-cellulo ubiquitination assay, followed by trypsin digestion to generate tryptic peptides. A second round of purification was then performed to enrich for KAP1-ubiquitinated peptides using GG remnant (K-ɛ-GG) immunoaffinity. After elution and desalting, the samples were subjected to LC-MS/MS. As with the other approaches, KAP1 samples submitted for MS analysis were validated for SMURF2-mediated ubiquitination (Fig. 1F).

Using these approaches, we identified three lysine residues on KAP1 that are ubiquitinated by SMURF2 (out of the 42 lysines present in its sequence): K254, K319 and K779 (Fig. 1G; Table S1). Notably, these residues have also been reported as potential ubiquitination sites of KAP1 in high-throughput MS analyses examining proteome-wide protein modifications (Table S2 and PhosphoSitePlus: https://www.phosphosite.org)^35^, with our findings suggesting that they are sites of SMURF2-mediated ubiquitination.

### Inactivating all three KAP1 residues (K254, K319, K779) is required to impair its ubiquitination by SMURF2

Following the identification of these KAP1 lysines as potential SMURF2 ubiquitination sites, we sought to determine whether they serve as bona fide ubiquitin acceptors. To this end, we mutated each of these lysines to arginine (K-to-R mutations) using site-directed mutagenesis, as well as generated double-mutant combinations involving these residues: K254R/K319R, K254R/K779R and K319R/K779R. We then examined the ability of SMURF2 to ubiquitinate these KAP1 mutants using an in-cellulo ubiquitination assay. Briefly, KAP1 wild-type (KAP1-WT) and its mutants were expressed in HEK-293T cells as FLAG-tagged proteins, either with or without MYC-SMURF2 and HA-tagged ubiquitin. After affinity purification with FLAG resin and SDS-PAGE, the mutants were analyzed for SMURF2-mediated ubiquitination by immunoblotting with an anti-HA (ubiquitin) antibody, comparing the results to KAP1-WT. While the data showed no significant reduction in ubiquitination levels for any single or double KAP1 mutant (Fig. 2A, B), it suggested that inactivating all three lysine sites may be necessary to affect SMURF2-mediated ubiquitination of KAP1. To examine this possibility, we generated a KAP1 triple mutant (KAP1-TM), incorporating mutations at all three lysine residues (K254R/K319R/K779R), and repeated an in-cellulo ubiquitination assay as described above. The results (Fig. 2C, D) revealed a profound reduction in SMURF2-mediated ubiquitination of KAP1-TM compared to its WT form, indicating that alterations at all three sites are required to impair SMURF2-mediated ubiquitination of KAP1.

**Figure 2.**
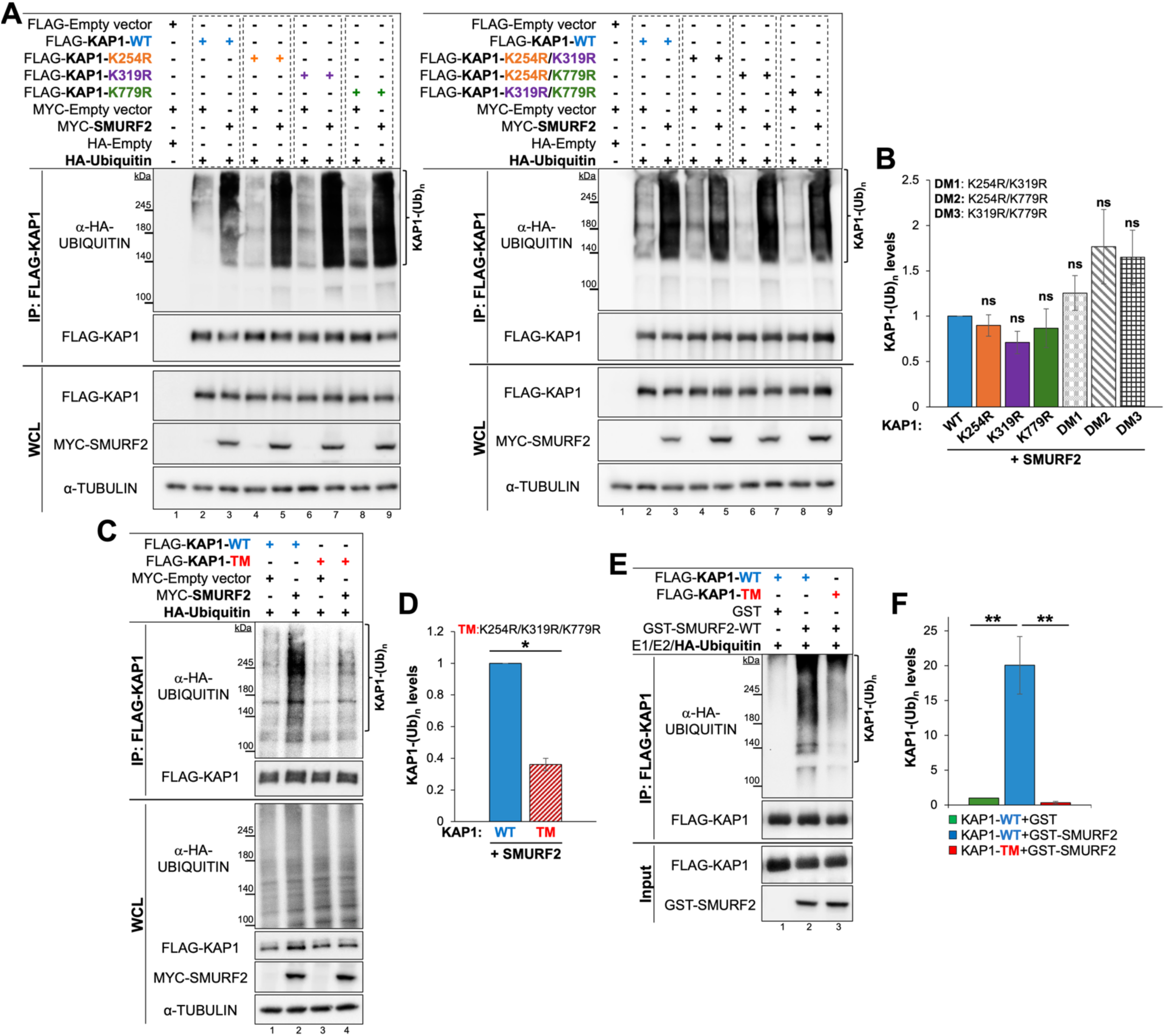
Inactivation of all three KAP1 residues (K254, K319, K779) is required to impair SMURF2-mediated ubiquitination of KAP1. **(A)** Western blot analysis of in-cellulo ubiquitination assays with single mutants (left panel) and double mutant combinations (right panel) of KAP1 (K-to-R mutations). **(B)** Quantification of the data shown in (A) from three independent experiments. Data are presented as mean ± SEM. ns, not significant. **(C)** Western blot analysis of in-cellulo ubiquitination assay showing SMURF2-mediated ubiquitination of KAP1-WT versus its mutant TM form. **(D)** Quantification of the data shown in panel (C) from three independent experiments. Data are mean ± SEM. \**P* < 0.05. **(E)** Western blot analysis of an in vitro ubiquitination assay, performed with purified KAP1 (WT or TM) and components of the ubiquitination cascade: E1 activating enzyme, E2 conjugating enzyme, HA-Ubiquitin, and E3 ubiquitin ligase SMURF2. **(F)** Quantification of the data shown in (E) from three independent experiments. Data are mean ± SEM. \*\**P* < 0.01.

To further substantiate these results, we examined SMURF2-mediated ubiquitination of purified KAP1-TM versus its wild-type form using an in vitro ubiquitination assay. Consistent with in-cellulo findings, the ubiquitination of KAP1-TM by SMURF2 was dramatically reduced compared to KAP1-WT (Fig. 2E, lane 3 vs. 2, and Fig. 2F).

Together, these data demonstrate that inactivation of all three KAP1 ubiquitination sites (K254, K319 and K779) is necessary to impair SMURF2-mediated ubiquitination.

### SMURF2 facilitates the recruitment of the deubiquitinase USP7 to KAP1 to regulate its ubiquitination-deubiquitination cycles through the K254/K319/K779 residues

Ubiquitination is a dynamic, tightly regulated process that depends on the coordinated activities of both E3 ubiquitin ligases and deubiquitinating enzymes (DUBs), which govern the spatio-temporal nature of ubiquitination and affect its outcomes.

In a previous study examining the impact of SMURF2 on the KAP1 interactome, we noted the deubiquitinase USP7/HAUSP as a potential KAP1 interactor^34^. Moreover, USP7 was also detected in the SMURF2 interactome^36^, suggesting that SMURF2 may act in concert with USP7 to regulate the ubiquitination of KAP1.

To test this hypothesis, we first performed a reciprocal co-immunoprecipitation (co-IP) analysis using recombinant FLAG-KAP1, MYC-SMURF2, and endogenous USP7, detecting complex formation between all three proteins (Fig. 3A). The results also suggested that increased expression of SMURF2 may enhance the interaction between KAP1 and USP7.

**Figure 3.**
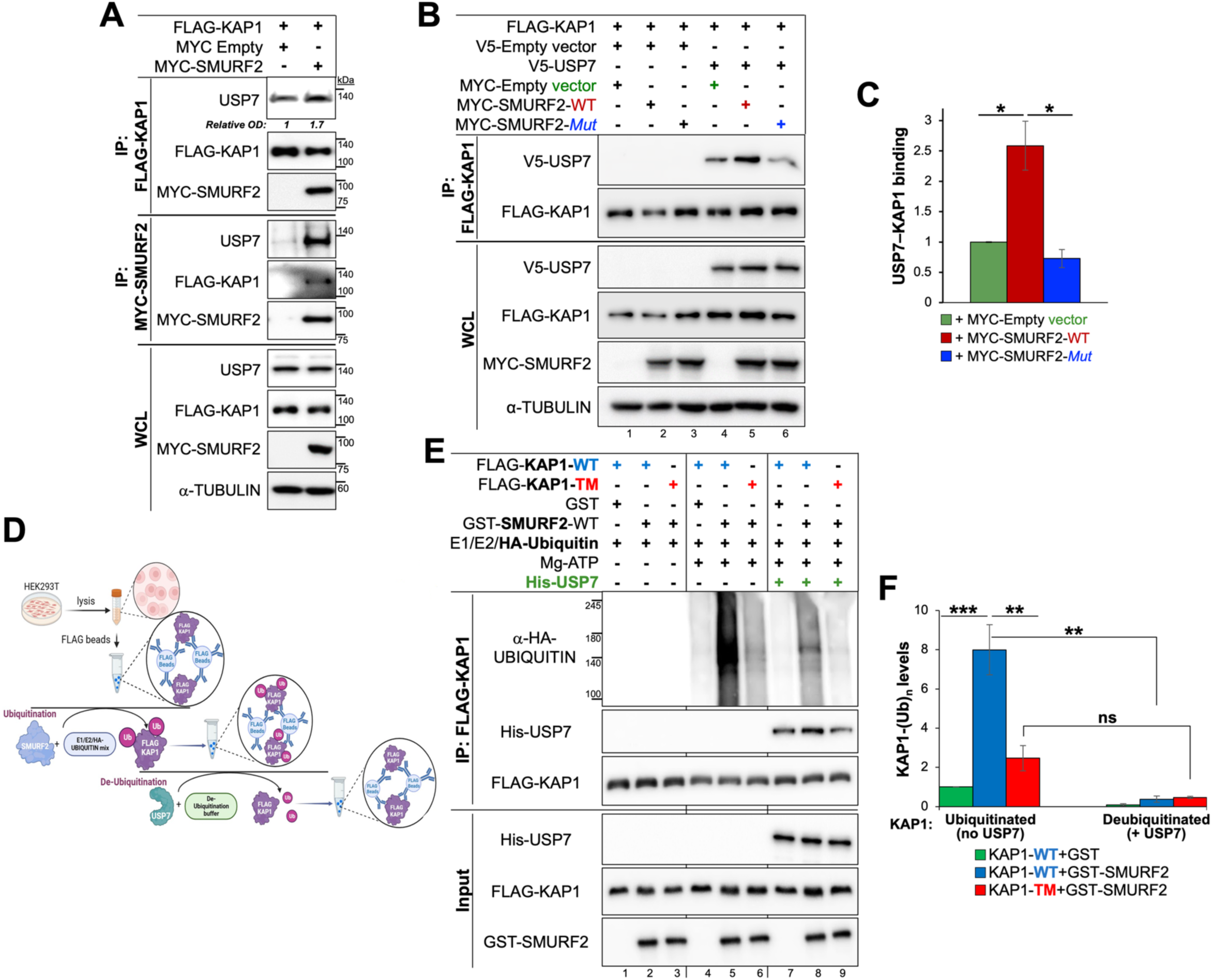
SMURF2 promotes the recruitment of the deubiquitinase USP7 to KAP1, regulating its ubiquitination-deubiquitination cycle via the identified lysine residues. **(A)** Reciprocal co-immunoprecipitation (co-IP) analysis showing interactions between endogenous USP7, FLAG-KAP1, and MYC-SMURF2 in HEK-293T cells. **(B)** Western blot analysis of FLAG-KAP1 immunoprecipitated from HEK-293T cells, with or without V5-USP7. Note the enhanced binding of USP7 to KAP1 facilitated by catalytically active SMURF2-WT, but not by its E3 ligase-dead SMURF2^C716G^ mutant (lane 6 vs. lane 5). **(C)** Quantification of the data shown in panel (B) from three independent experiments. Data are mean ± SEM. \**P* < 0.05. **(D)** Schematic diagram of the ubiquitination-deubiquitination assay workflow used to identify USP7 as KAP1’s deubiquitinating enzyme following SMURF2-mediated ubiquitination. **(E)** Western blot analysis of SMURF2/USP7-regulated ubiquitination and deubiquitination of KAP1. Samples without Mg-ATP were included as additional controls to validate the specificity of the reaction (lanes 1-3 vs. lanes 4-9). Note the protein complex formation between purified USP7 and KAP1 (lanes 7-9). **(F)** Quantification of the data shown in panel (E) from three independent experiments. Data are mean ± SEM. \*\**P* < 0.01; \*\*\**P* < 0.001. ns, not significant.

To further validate this finding, we conducted additional co-IP experiments using FLAG-KAP1, V5-tagged USP7 (V5-USP7), and MYC-SMURF2 (either WT or Mut) expressed in HEK-293T cells, with empty V5 and MYC vectors serving as additional controls. The results confirmed complex formation between KAP1 and USP7 and showed that catalytically active SMURF2 markedly enhances this interaction (Fig. 3B, lane 5 vs. lanes 4 and 6, and Fig. 3C), suggesting that SMURF2 promotes USP7 recruitment to KAP1.

Next, to determine whether USP7 functions as the enzyme responsible for deubiquitinating KAP1, we conducted an in vitro ubiquitination assay followed by KAP1 deubiquitination analysis, as outlined in Fig. 3D and detailed in the *Materials and Methods* section. Briefly, FLAG-KAP1-WT and KAP1-TM, purified and immobilized on beads, were incubated with E1, E2, GST-SMURF2, and HA-ubiquitin for the ubiquitination reaction. The samples were then washed, resuspended in deubiquitination buffer, and incubated with purified His-USP7. After the deubiquitination reaction, the samples were washed again, FLAG-KAP1 was eluted, and the proteins were analyzed by Western blotting (Fig. 3E).

Consistent with our previous findings, SMURF2 significantly increased the ubiquitination of KAP1-WT, an effect that was markedly reduced in the KAP1-TM samples (Fig. 3E, lane 5 vs. lanes 6 and 4; Fig. 3F). Addition of USP7 to ubiquitinated KAP1-WT efficiently removed its SMURF2-mediated ubiquitination (Fig. 3E, lane 8 vs. lane 5; Fig. 3F), whereas KAP1-TM remained largely resistant to deubiquitination by USP7 (Fig. 3F).

Collectively, these findings reveal that SMURF2 collaborates with the deubiquitinase USP7, promoting its interaction with KAP1 to regulate its ubiquitination-deubiquitination cycle via the K254/K319/K779 residues.

### SMURF2/USP7-refractory KAP1 has impaired SUMO E3 ligase activity

KAP1 regulates chromatin structure and gene expression by interacting with various transcriptional co-repressors and chromatin-modifying enzymes. This function of KAP1 depends on its intrinsic SUMO E3 ligase activity, which is essential for recruiting chromatin remodelers to the genome. To investigate the consequences of disrupting SMURF2/USP7-mediated regulation of KAP1, we first examined its impact on KAP1’s SUMO E3 ligase activity. To this end, we conducted KAP1 auto-SUMOylation assays using purified components of the SUMOylation cascade: SUMO-activating enzyme E1, SUMO-conjugating enzyme Ubc9, SUMO1, and FLAG-tagged KAP1 (WT or KAP1-TM) purified from HEK-293T cells. After the SUMOylation reaction, KAP1 was eluted from the beads and analyzed for auto-SUMOylation by Western blotting using an anti-SUMO1 antibody. Purified Ran GTPase-activating protein 1 (RanGAP1), with or without Mg-ATP, was used as the positive or negative control for the reaction, respectively (Fig. 4A, lanes 1 and 2). As previously reported (Ivanov et al., 2007; Zeng et al., 2008), KAP1-WT exhibits prominent auto-SUMOylation activity (Fig. 4A, lane 4). In contrast, KAP1-TM showed a significant reduction in its ability to auto-SUMOylate compared to its WT counterpart (Fig. 4A, lane 5 vs. lane 4; and Fig. 4B).

**Figure 4.**
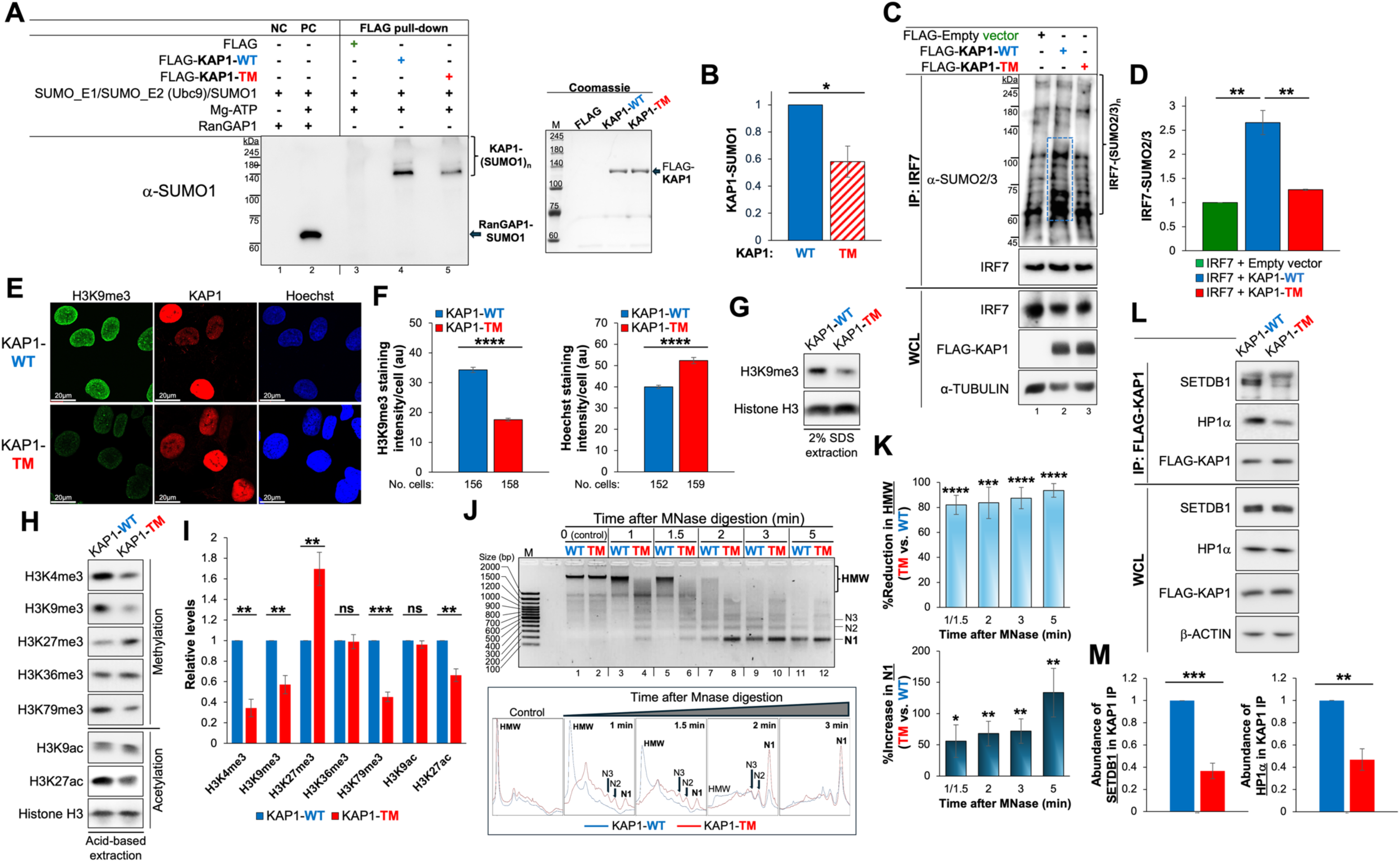
KAP1-TM exhibits impaired SUMO E3 ligase activity, leading to global alterations in the epigenetic landscape and chromatin structure. **(A)** Western blot analysis of KAP1 auto-SUMOylation using purified FLAG-KAP1-WT and FLAG-KAP1-TM in the presence of purified components of the SUMOylation machinery. FLAG-KAP1 was pulled down from the reaction using anti-FLAG beads and probed with anti-SUMO1 antibody. Purified RanGAP1, with or without Mg-ATP, was used as a positive or negative control, respectively (lanes 1 and 2). PC, positive control; NC, negative control. Coomassie staining (bottom panel) confirms the purity and equal loading of FLAG–KAP1-WT and FLAG-KAP1-TM proteins used in the reaction. **(B)** Quantification of KAP1-WT and KAP1-TM auto-SUMOylation from three independent experiments. Data are mean ± SEM. \**P* < 0.05. **(C)** In-cellulo SUMOylation assay of endogenous IRF7 immunoprecipitated from HEK-293T cells. **(D)** Quantification of the data presented in panel (C) from three independent experiments. Data are mean ± SEM. \*\**P* < 0.01. **(E)** Confocal images of KAP1-replaced U2OS cells showing the effects of KAP-TM expression on H3K9me3 levels and chromatin compaction. Cell nuclei were stained with Hoechst 33258, and FLAG-KAP1 was detected by anti-FLAG immunofluorescence. Scale bars, 20 μm. **(F)** Quantification of H3K9me3 (left panel) and Hoechst 33258 staining intensity (right panel) in KAP1-replaced U2OS cells from three independent experiments. The number of cells used for quantification is shown in the figure. Data are mean ± SEM. \*\*\*\**P* < 0.0001. **(G)** Western blot analysis of H3K9me3 levels in KAP1-WT and KAP1-TM U2OS cells. Cells were lysed in 2% SDS, followed by immediate sample boiling, as detailed in the Materials and Methods. **(H)** Western blot analysis of histone H3 methylation and acetylation levels in KAP1-WT and KAP1-TM cells. Histone were purified by acid-based extraction. **(I)** Quantification of the data presented in panel (H) from three separate experiments. Data are mean ± SEM. \*\**P* < 0.01. ns, not significant. **(J)** MNase digestion assay. N1, N2, and N3 indicate DNA fragments corresponding to mono-, di- and tri-nucleosomes, respectively. HMW, high molecular weight DNA. M, DNA marker. The bottom panel shows densitometric profiles of untreated and MNase-digested chromatin. **(K)** Quantification of four independent MNase digestion experiments. The left panel shows the percentage reduction in high molecular weight (HMW) DNA in KAP1-TM versus WT cells, while the right panel shows the percentage increase in N1 mononucleosome intensity. Data are presented as mean ± SEM. \**P* < 0.05; \*\**P* < 0.01; \*\*\**P* < 0.001; \*\*\*\**P* < 0.0001. **(L)** Co-IP analysis showing interactions between KAP1 (WT or TM) and endogenous SETDB1 and HP1α in KAP1-replaced U2OS cells. **(M)** Quantification of the data shown in (L) from three different experiments. Data are mean ± SEM. \*\**P* < 0.01; \*\*\**P* < 0.001.

To further substantiate our findings on the compromised catalytic activity of KAP1-TM, we assessed its ability to SUMOylate other cellular substrates, particularly interferon regulatory factor 7 (IRF7)−a key regulator of type I interferon-dependent immune responses and the first identified transcription factor SUMOylated by KAP1. To this end, we conducted in-cellulo SUMOylation assays, comparing the ability of KAP1-TM versus KAP1-WT to SUMOylate endogenously expressed IRF7. Briefly, IRF7 was immunoprecipitated from HEK-293T cells ectopically expressing either KAP1-WT or KAP1-TM, followed by Western blot analysis of its SUMOylation by endogenous SUMO2/3 proteins. The results (Fig. 4C, D) show that while KAP1-WT efficiently promotes the SUMOylation of IRF7, as evidenced by a significant increase in IRF7 SUMOylation in KAP1-WT-expressing cells, KAP1-TM failed to induce this effect, with SUMOylation levels comparable to the cells transduced with an empty vector (Fig. 4C, lane 3 vs. lanes 1, 2; Fig. 4D).

Cumulatively, these findings demonstrate that disrupting the SMURF2-USP7-mediated regulation of KAP1 impairs its SUMO E3 ligase activity.

### The dysregulated SMURF2/USP7-KAP1-SUMOylation axis leads to global alterations in the epigenetic landscape and chromatin structure

Impaired SUMO E3 ligase activity of KAP1 suggests that its disrupted regulation may cause widespread changes in chromatin, leading to functional alterations. To investigate these effects, we generated KAP1-replaced cells on the background of U2OS cells, a widely used model for studying nuclear organization and functions. In these cells, we stably replaced the endogenous KAP1 with either its WT or TM form (Fig. S1). This protein-replacement approach offers several advantages. First, it enables the assessment of SMURF2/USP7 activities exerted specifically through KAP1, avoiding interference from other cellular substrates of these enzymes. Second, it eliminates the interference of endogenous KAP1 in the experimental system, providing a more controlled environment for analysis. Third, it facilitates the generation of stable cell models for subsequent investigations, including in vitro and in vivo analyses.

Using these cells, we first examined histone H3 trimethylation at lysine 9 (H3K9me3), an epigenetic hallmark of constitutive heterochromatin. This modification is regulated by histone methyltransferases such as SETDB1, which are recruited to chromatin by auto-SUMOylated KAP1, facilitating the formation and/or maintenance of heterochromatin structure^15,17,37^.

Immunostaining and western blot analyses revealed significantly reduced levels of H3K9me3 in KAP1-TM cells (Fig. 4E, F, left panels; and Fig. 4G-I). Notably, alongside the reduction in H3K9me3, we also observed substantial decreases in several other histone modifications, including trimethylation of H3 at K4 (H3K4me3) and K79 (H3K79me3), as well as acetylation at K27 (H3K27ac). In contrast, levels of H3 trimethylation at K27 (H3K27me3) were significantly elevated (Fig. 4H, I), while trimethylation at K36 (H3K36me3) and acetylation at K9 (H3K9ac) remained unchanged. These findings suggest that SMURF2/USP7-refractory KAP1 is unable to effectively support the epigenetic landscape associated with heterochromatin, leading to alterations in chromatin states.

Supporting this conclusion, staining of KAP1-replaced cells with Hoechst 33258, a DNA dye that binds specifically to the minor groove and highlights relaxed chromatin structure^38–40^, showed markedly increased fluorescence intensity in KAP1-TM cells, indicating a more relaxed chromatin conformation (Fig. 4E, F, right panels).

Furthermore, to directly assess the impact of KAP1-TM on chromatin architecture, we performed a micrococcal nuclease (MNase) sensitivity assay, which evaluates chromatin compaction in its native state by measuring DNA accessibility within intact nuclei^40,41^. Consistent with our other findings indicating chromatin decompaction in KAP1-TM cells, these cells exhibited significantly increased chromatin digestion, indicative of more loosely compacted chromatin (Fig. 4J, K).

Finally, to explore the molecular mechanisms underlying these changes, we examined the ability of KAP1-TM to interact with two major KAP1-binding partners involved in heterochromatin formation and maintenance: SETDB1 and HP1α. Our results show that complex formation between KAP1-TM and these proteins was profoundly diminished (Fig. 4L, M).

### Impaired regulation of KAP1 leads to altered gene expression, affecting multiple cellular pathways

Given the global chromatin changes in KAP1-TM cells, we hypothesized that dysregulation of KAP1 could result in significant changes in gene expression, as structural changes in chromatin can directly affect the accessibility of the transcriptional machinery. To test this hypothesis, we performed RNA-Seq-based gene expression analysis, comparing KAP1-TM to KAP1-WT cells. Overall, we detected in these cells the expression of 27,288 transcripts, including both protein-coding and non-coding genes. After applying additional thresholds (i.e., changes in gene expression > 25%, FDR < 1%, and *P*-value < 0.05), we identified 298 genes with significantly altered expression. Of these, 116 genes exhibited elevated expression, while 182 showed decreased expression in KAP1-TM compared to WT cells (Fig. 5A, B; Table S3). Notably, in addition to the dysregulated expression of protein-coding genes, we identified significant changes in the expression of non-coding RNAs, particularly lncRNAs, with 33 downregulated and 26 upregulated transcripts (Fig. S2A).

**Figure 5.**
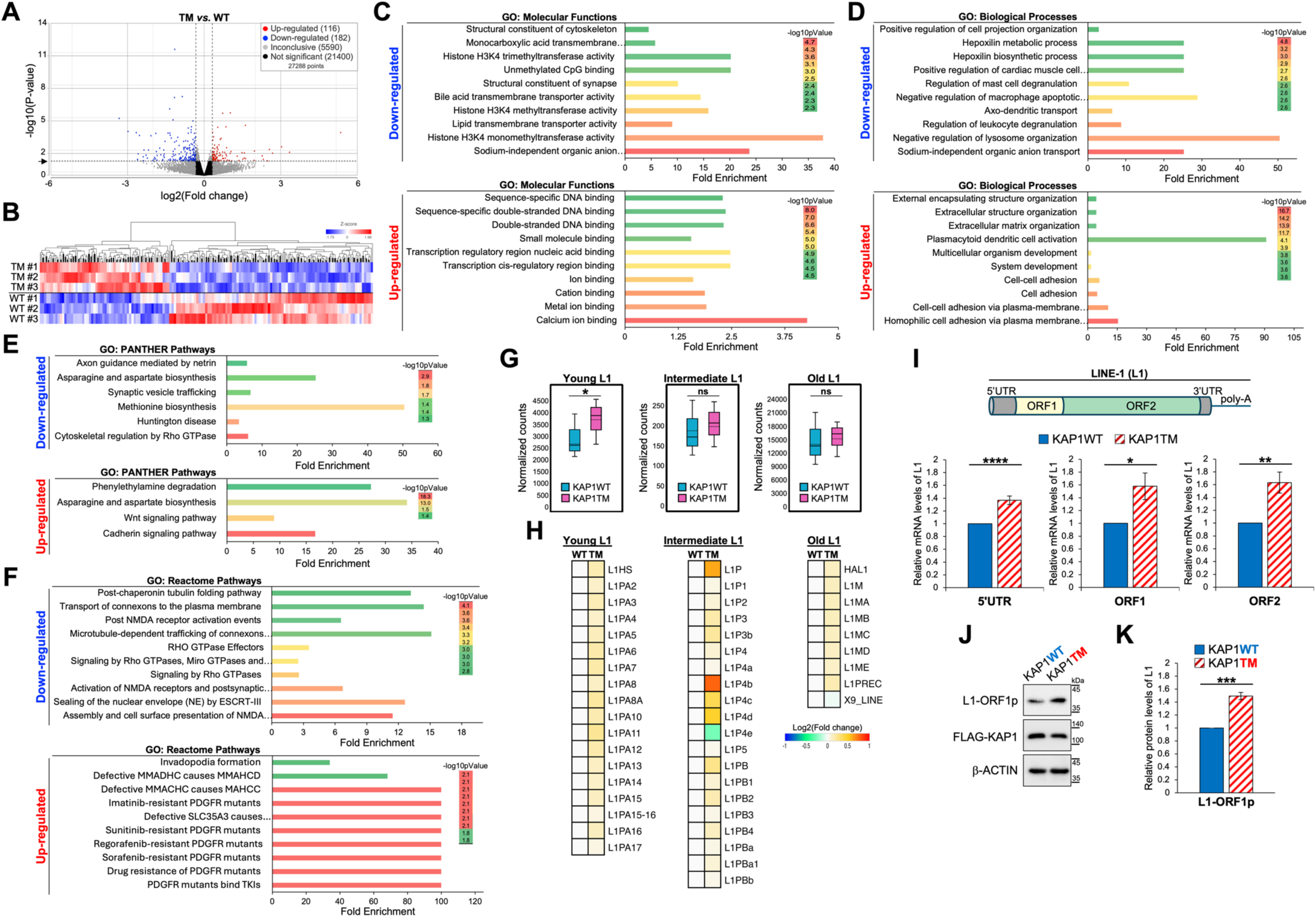
KAP1-TM disrupts gene expression programs and elevates LINE-1 retrotransposon expression. **(A)** Volcano plot showing RNA-Seq results from U2OS cells expressing either KAP1-WT or KAP1-TM, each with three biological replicates. Differential gene expression was defined by a threshold of >25% change, FDR < 1%, and *P* < 0.05. Genes upregulated and downregulated in KAP1-TM relative to WT controls are highlighted in red and blue, respectively. **(B)** Hierarchical clustering heatmap of differentially expressed genes in KAP1-TM vs. WT cells. **(C-F)** GO enrichment analysis of molecular functions, biological processes, and cellular pathways up- and down-regulated in KAP1-TM versus WT cells. The fold enrichment for each GO category is plotted on the X-axis. The effects on cellular pathways were assessed using two complementary approaches: GO:PANTHER (panel E) and GO:Reactome pathway analysis (panel F). The top ten categories with the most significant *p*-values are shown for each GO term. The full datasets for each GO category are provided in Supplemental Tables S6-S8. **(G)** Box plots showing the effect of KAP1-TM on the expression of young, intermediate and old LINE-1 retroelements. The analysis was conducted using the Plotly Python package (https://plotly.com/python). Data are mean ± SD of three biological replicates. \**P* < 0.05. **(H)** Heatmaps of the individual LINE-1 elements expressed in KAP1-WT and KAP1-TM cells, showing the average values of three biological replicates for each LINE-1 element. The heatmaps were generated using Morpheus software (https://software.broadinstitute.org/morpheus). **(I)** RT-qPCR analysis showing the effect of KAP1-TM on LINE-1 expression using three distinct sets of primers targeting different LINE-1 regions: 5’UTR, ORF1, and ORF2. Data are mean ± SEM of three biological replicates. \**P* < 0.05; \*\**P* < 0.01; \*\*\*\**P* < 0.0001. **(J)** Western blot analysis of LINE-1 ORF1 protein (L1-ORF1p) expression in KAP1-replaced cells. **(K)** Quantification of data shown in panel (J) from four biological replicates. Data are mean ± SEM. \*\*\**P* < 0.001.

Cross-referencing our RNA-Seq findings with available KAP1 ChIP-Seq data^42^ revealed that nearly 44% of the genes differentially expressed in KAP1-TM versus WT cells (130 out of 298 genes; 43.6%) contain KAP1 binding sites (Fig. S2B, C; Table S4), suggesting that KAP1 may directly regulate their expression.

To validate the robustness of our RNA-Seq findings, we selected five different genes whose expression was either upregulated or downregulated in KAP1-TM compared to WT cells and analyzed their expression using quantitative real-time PCR (RT-qPCR). The selection was also based on two additional criteria: (i) the presence of KAP1 binding sites in the selected genes, determined through KAP1 ChIP-Seq data analysis, and (ii) consistent changes in gene expression direction in both KAP1-TM and KAP1 knockdown (siKAP1) cells (Tables S4, S5; and ref.^43^). The RT-qPCR results confirmed the gene expression changes detected in our RNA-Seq analysis (Fig. S2D), further supporting the reliability of the RNA-Seq findings.

Subsequent Gene Ontology (GO) analysis revealed that the genes differentially expressed between KAP1-WT and KAP1-TM cells are implicated in various molecular functions and biological processes (Fig. 5C and D; Tables S6 and S7). For instance, genes associated with histone methyltransferase activity were significantly downregulated in KAP1-TM cells, while genes involved in DNA binding and metal/ion binding were upregulated.

Biological processes downregulated in KAP1-TM cells included positive regulation of cell projection organization, protein oligomerization, negative regulation of lysosome organization, and sodium-independent organic anion transport, among 358 other downregulated processes (Fig. 5D; Table S7). Additionally, processes related to cell fate specification, differentiation, antisense RNA metabolism, and snoRNA catabolism were also downregulated in these cells. In contrast, biological processes enriched in KAP1-TM cells included extracellular structure organization, cell adhesion, morphogenesis, organelle organization, cellular responses to stress and stimuli, RNA processing, regulation of piRNA transcription, and metabolic processes, with a total of 205 significantly upregulated processes (Fig. 5D; Table S7).

Furthermore, GO pathway analysis showed that genes downregulated in KAP1-TM cells are involved in key cellular pathways, including signal transduction (e.g., Rho GTPase, AMPK, AKT, and PKR signaling), regulation of TP53 expression and degradation, tubulin folding, gap junction regulation, amino acid biosynthesis, mTORC1 regulation, and fatty acid metabolism, among others (Fig. 5E, F, upper panels; Table S8). In contrast, the upregulated pathways in KAP1-TM cells included WNT and cadherin signaling, asparagine and aspartate biosynthesis, invadopodia formation, as well as pathways associated with anticancer drug response (Fig. 5E, F, bottom panels; Table S8).

Taken together, these results demonstrate that dysregulation of KAP1 leads to significant changes in gene expression, affecting a wide range of biological processes and diverse cellular pathways.

### Dysfunctional SMURF2/USP7-KAP1-SUMOylation axis leads to elevated LINE-1 retrotransposon expression

KAP1 plays a crucial role in preserving genomic integrity by silencing transposable elements, particularly LINE-1, a highly abundant and active family of retrotransposons. This silencing is essential for maintaining chromatin structure and preventing genomic instability, a hallmark of age-related disorders such as cancer^44–46^.

To investigate whether disruption of SMURF2/USP7-mediated regulation of KAP1 affects LINE-1 expression, we employed several approaches. First, using our RNA-seq data, we compared LINE-1 expression levels in KAP1-TM and WT cells, categorizing LINE-1 elements by their evolutionary age: young (human-specific), intermediate (primate-specific), and old (mammalian-specific)^47,48^. As shown in Figs. 5G and 5H, KAP1-TM cells exhibited significantly elevated expression of LINE-1 (L1) transcripts, particularly young L1s. Similarly, KAP1 knockdown cells also showed a significant increase in young L1 expression, though intermediate L1s were also upregulated in these cells (Fig. S2E, F). The expression of old, mammalian-specific L1s did not show statistically significant changes in either KAP1-TM or KAP1 knockdown cells.

The elevated expression of L1 in KAP1-TM cells was further validated by RT-qPCR analysis using three distinct sets of primers targeting different LINE-1 regions: 5’UTR, open reading frame 1 (ORF1), and open reading frame 2 (ORF2). The results (Fig. 5I) confirmed a significant upregulation of L1 expression in KAP1-TM cells, with approximately a 50% increase in LINE-1 expression, consistent with our RNA-Seq results.

Additionally, Western blot analysis using an anti-ORF1p-specific antibody, which detects the expression of a nucleic acid-binding protein, ORF1p, essential for LINE-1 retrotransposition^49^, revealed significantly elevated expression of L1 in KAP1-TM cells (Fig. 5J, K), further supporting our RNA-Seq and RT-qPCR findings.

Collectively, these results demonstrate that disruption of the SMURF2/USP7-KAP1-SUMOylation axis leads to derepression of LINE-1 elements in KAP1-TM cells. This primarily affects young, transcriptionally active L1s, which tend to be located closer to genes than older L1s^44,50^, and are particularly susceptible to chromatin structure changes observed in KAP1-TM cells.

### Impaired KAP1 regulation disrupts protein-protein interactions (PPIs), affecting multiple cellular processes and functions

KAP1 regulates chromatin structure and gene expression by acting as a scaffold for diverse PPIs with various cellular factors. Disruptions in KAP1 function, observed in KAP1-TM cells, suggest that its protein interactions may be significantly altered, as exemplified by markedly reduced interactions of KAP1-TM with SETDB1 and HP1α (Fig. 4L, M). To assess how impaired KAP1 regulation impacts its PPIs on a global scale, we performed MS-based proteomic analysis comparing the KAP1 interactome in KAP1-TM versus WT cells. Briefly, FLAG-tagged KAP1 (WT and TM) was expressed and affinity-purified from HEK-293T cells, chosen for their ability to produce high yields of recombinant proteins, which is essential for comprehensive PPI mapping. After extensive washing and elution of KAP1-containing protein complexes, the samples were subjected to LC-MS/MS analysis, as detailed in *Materials and Methods*. The efficiency of KAP1 pull-down from WT and TM cells was validated by Western blot analysis and Coomassie gel staining (Fig. S3A).

Using this approach, we identified 808 proteins interacting with KAP1-WT and 752 proteins in the KAP1-TM interactome (Fig. 6A). Among these, 629 proteins (∼77%) were shared between the two interactomes, indicating that a substantial portion of KAP1 PPIs is conserved across the WT and TM samples. Nonetheless, 179 and 123 proteins were found to be specifically associated with KAP1-WT and KAP1-TM, respectively.

**Figure 6.**
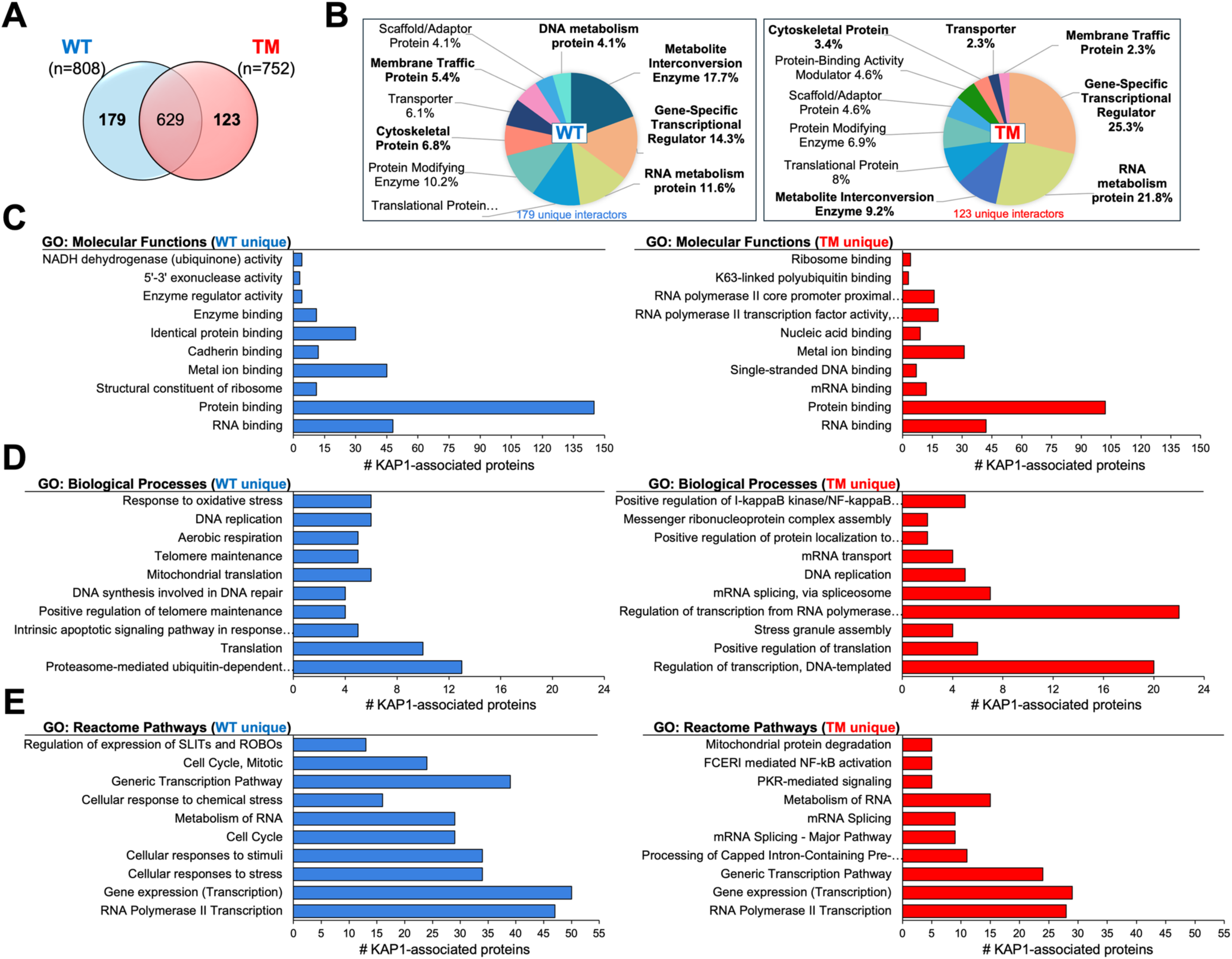
Impaired regulation of KAP1 disrupts its PPIs, affecting multiple cellular processes and pathways. **(A)** Venn diagram showing the total number of KAP1-associated proteins identified by LC-MS/MS in cells expressing either KAP1-WT or KAP1-TM. All identified peptides were filtered for high confidence, top rank, mass accuracy, and a minimum coverage of two peptides. **(B)** Protein classes of KAP1 unique interactors determined using the PANTHER classification system. The top ten categories with the highest number of protein interactors are shown. Full data are provided in Supplemental Table S9. Protein classes highlighted in bold indicate >50% changes in abundance between KAP1-WT and KAP1-TM samples. **(C-E)** GO enrichment analysis of molecular functions, biological processes, and cellular pathways in KAP1-TM versus WT cells, based on their unique protein interactors. The number of KAP1-associated proteins for each GO category in the KAP1-WT and TM interactomes is plotted on the X-axis. The top ten categories with the most significant *p*-values are shown. Full datasets are provided in Supplemental Tables S10-S12. Note, although some GO terms were shared between the two interactomes, the number of KAP1-associated proteins was markedly reduced in the KAP1-TM interactome.

Functional classification of these unique interactors using the PANTHER classification system showed that although they belong to similar protein classes (e.g., DNA and RNA metabolism, chromatin binding, gene-specific transcriptional regulation, translational proteins, and metabolite interconversion enzymes), their composition and protein abundance within the classes were significantly altered (Fig. 6B). Furthermore, certain protein classes, such as extracellular matrix proteins, intercellular signaling molecules, and proteins associated with defense and immunity, were detected only among the KAP1-TM interactors (unique interactors group; Fig. 6B; Table S9).

PPI maps, generated using the STING tool, further demonstrated distinct associations of KAP1’s unique interactors with various functional networks. For KAP1-WT, these networks included DNA replication and repair, negative regulation of transcription (DNA-templated), translation, and cellular respiration. In contrast, KAP1-TM interactors enriched functional networks related to negative regulation of the G1/S transition, mitotic cell cycle, mRNA splicing via the spliceosome, and positive regulation of translation (Fig. S3B).

GO enrichment analysis of molecular functions revealed further differences between the two KAP1 interactomes. Specifically, 21 out of the 27 molecular functions associated with KAP1-WT were absent in the KAP1-TM interactome. These included identical protein and enzyme binding; unfolded protein binding; binding of damaged DNA; microtubule binding; and proteasome binding, among others (Fig. 6C; Table S10). In contrast, the KAP1-TM interactome was enriched for single-stranded DNA binding, nucleic acid binding, sequence-specific DNA binding associated with RNA polymerase II transcriptional activity, and K63-linked polyubiquitin binding. Functions shared by both interactomes included RNA and protein binding, metal ion binding, ribosome binding, and sequence-specific DNA binding at the RNA polymerase II core promoter proximal region (Fig. 6C; Table S10).

Further analysis of biological processes associated with the unique interactors in the KAP1-WT and TM interactomes indicated that while some processes were common to both (e.g., regulation of DNA replication, transcription, and mRNA transport), the majority (47 out of 51) were exclusively enriched in the KAP1-WT interactome (Fig. 6D; Table S11). These processes included proteasome-mediated ubiquitin-dependent protein catabolic process, translation, DNA repair and recombination, telomere maintenance, cellular respiration, response to oxidative stress, protein folding, cell-cell adhesion, and regulation of cell growth and proliferation, among others. In contrast, processes enriched in the KAP1-TM interactome comprised negative regulation of transcription from the RNA polymerase II promoter, regulation of histone phosphorylation, mRNA polyadenylation, positive regulation of translation, stress granule assembly, and NF-kappa-B signaling (Fig. 6D; Table S11).

In line with these findings, GO pathway analysis revealed prominent alterations across various cellular pathways, underscoring the impact of KAP1-TM on cellular functions (Fig. 6E; Table S12). Remarkably, out of 276 pathways linked to KAP1-WT, only 16 were detected in KAP1-TM cells. The shared pathways included transcription regulation, RNA processing and splicing, metabolism of RNA and proteins, mitochondrial protein degradation, Rho GTPase, FCERI and Hedgehog signaling, as well as the regulation of the mitotic G1 phase and G1/S transition. Conversely, the pathways specifically associated with KAP1-TM included JNK and p38 MAPK activation, PKR-mediated signaling, interleukin-17 signaling, Toll-like receptor signaling, NOD1/2 signaling, antiviral mechanism by IFN-stimulated genes, and the post-chaperonin tubulin folding pathway (Fig. 6E; Table S12).

Finally, to gain a comprehensive understanding of the effects associated with impaired KAP1 regulation through its PPIs, we also analyzed KAP1 interactions within the shared protein group (629 common interactors), using a two-fold change in protein abundance as the threshold for identifying differential interactions. This analysis identified 157 proteins specifically associated with KAP1-WT, compared to only 36 interactors in the KAP1-TM group, revealing substantial differences in protein classes and functional networks (Fig. S3C-E; Tables S13). Subsequent GO analyses indicated significant disparities between the KAP1-WT and KAP1-TM interactomes across all analyzed GO categories (Fig. S2F-H; Tables S14-S16). The most affected biological processes included the negative regulation of transcription from the RNA polymerase II promoter, regulation of mRNA and protein stability, protein folding, translation, post-translational modifications, ubiquitin-dependent proteasomal degradation, protein polyubiquitination and deubiquitination, regulation of the cellular stress response and response to DNA damage, WNT, NIK/NF-kappa-B, and MAPK signaling pathways, and cell cycle regulation, along with 72 other processes associated with KAP1-WT but absent in the KAP1-TM interactome (Fig. S3G; Table S15). Correspondingly, GO pathway analysis identified alterations in the cellular pathways related to metabolism, gene expression regulation, stress and stimulus responses, cell cycle regulation, programmed cell death, immune system regulation, viral infection pathways, and associated diseases (Fig. S3H; Table S16).

Together, these results demonstrate significant divergence in the molecular functions, biological processes, and cellular pathways associated with impaired KAP1, suggesting potential functional adaptations or disruptions in the cellular networks regulating critical cellular processes.

### Disruption of the SMURF2-KAP1 axis suppresses tumorigenicity

To further assess the impact of impaired KAP1 regulation, we examined the proliferation and tumorigenic potential of KAP1-replaced U2OS cells. We found that while short-term proliferation (24-96 hours) was comparable between KAP1-WT and TM cells (Fig. S4A), KAP1-TM cells exhibited markedly reduced colony-forming capacity over a longer period (12 days; Fig. 7A, B), suggesting that impaired KAP1 regulation may suppress cell tumorigenic potential. Consistent with these findings, bi-parametric FACS analysis following BrdU incorporation and PI staining revealed a slight but statistically significant reduction in the proportion of actively cycling KAP1-TM cells (Fig. S4B). This cell cycle defect likely contributes to the reduced colony size and diminished proliferative capacity of KAP1-TM cells over time, as observed in the colony formation assay.

**Figure 7.**
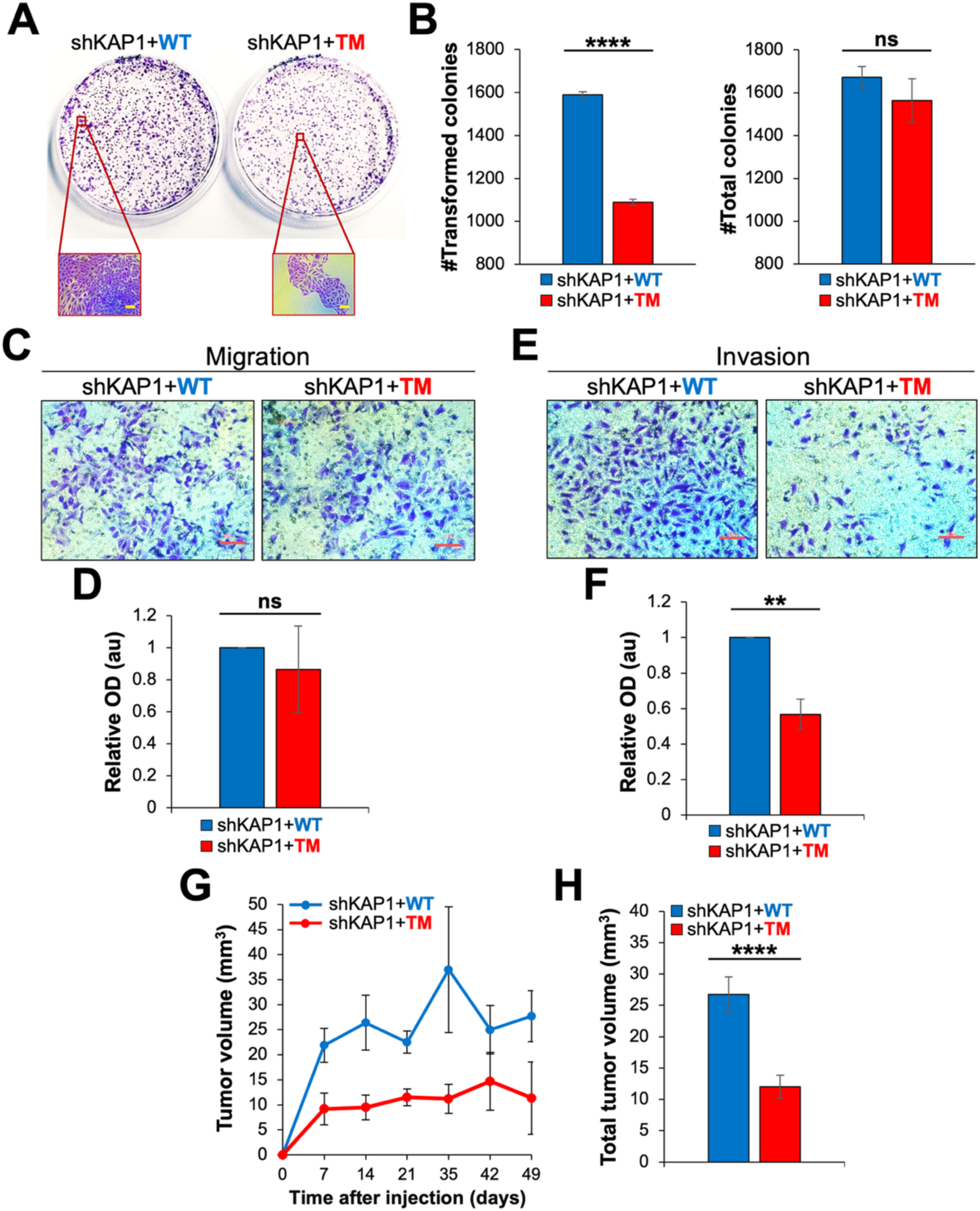
Disruption of the SMURF2/USP7-KAP1-SUMOylation axis suppresses cancer cell tumorigenicity and tumor growth in vivo. **(A)** Colony formation assay of KAP1-replaced U2OS cells. The images below show close-up views of the crystal violet-stained cell foci, illustrating the average size of colonies in KAP1-WT versus TM cells. Scale bar, 100 μm. **(B)** Quantification of the data obtained in colony formation assays. The left and right panels show the number (#) of transformed colonies and the total number of colonies, confirming equal cell plating, respectively. Data are mean ± SEM. \*\*\*\**P* < 0.0001. ns, not significant. **(C)** Representative images showing the transwell migration of KAP1-WT and TM U2OS cells. Scale bar, 100 μm. **(D)** Quantification of the results of cell migration assays obtained from three independent experiments. The stained filters were dissolved in 1% SDS and OD measurements were taken at 570 nm. Data are mean ± SEM. ns, not significant. **(E)** Representative images of U2OS cell invasion through the matrix (Matrigel) in a confined 3D environment (transwell invasion assay). Scale bar, 100 μm. **(F)** Quantification of the cell invasion data obtained from three independent experiments. Data are mean ± SEM. \*\**P* < 0.01. **(G)** Growth curves of tumors developed in NSG mice following subcutaneous injection of KAP1-WT or TM U2OS cells. Mice were injected with two million cells resuspended in a PBS/Matrigel solution. Tumor growth was monitored at the indicated time points using a caliper. Each group consisted of five mice. **(H)** The calculated total tumor volume in mice injected with KAP1-WT and TM U2OS cells. Data are mean ± SEM. \*\*\*\**P* < 0.0001.

Furthermore, the results showed that while the expression of KAP1-TM did not significantly affect the migratory capabilities of the cells (Fig. 7C, D), it substantially reduced their capacity to invade through the extracellular matrix in a confined 3D environment (Fig. 7E, F). Remarkably, similar outcomes were also observed in cells where the endogenous KAP1 was transiently replaced with the TM form, as well as in KAP1 knockdown cells (Fig. S4C-H). Reconstitution of KAP1 in these cells rescued this phenotype. Overall, the phenotype of KAP1 knockdown cells closely resembled the phenotype of cells expressing mutant KAP1, whether transiently or stably (Fig. 7A-F and Fig. S4A-H), indicating that the loss of KAP1 functionality, either through knockdown or mutation, produces similar outcomes, reducing the tumorigenic potential of the cells.

Finally, to assess the impact on tumorigenicity in vivo, we performed a xenograft tumor formation assay using NSG mice injected with either KAP1-WT or KAP1-TM U2OS cells. The results (Fig. 7G) show that tumors expressing KAP1-TM exhibited significantly slower growth rates and reduced tumor volumes compared to KAP1-WT control (Fig. 7G, H).

Taken together, our results elucidate the mechanism of KAP1 regulation and demonstrate its pivotal role in governing critical cellular processes, such as the maintenance of the epigenetic landscape, chromatin compaction, gene expression, retrotransposon activity, and protein-protein interactions. Our findings also highlight the significance of this mechanism in regulating cancer cell proliferation, invasion and tumor growth, suggesting that targeting this pathway could be a promising direction for cancer treatment.

## DISCUSSION

In this study, we identified a novel regulatory mechanism governing the activity of the core chromatin and transcriptional regulator KAP1. This mechanism operates at the intersection of ubiquitination dynamics and SUMO E3 ligase activity, influencing chromatin structure, gene expression and key cellular processes. We demonstrate that KAP1 undergoes non-proteolytic ubiquitination at three specific lysine residues (K254, K319 and K779), mediated by the E3 ubiquitin ligase SMURF2, which modulates KAP1’s SUMOylation activity. Our findings further show that KAP1 ubiquitination is tightly regulated by the coordinated action of SMURF2 and the deubiquitinase USP7, which is recruited to KAP1 by SMURF2 in a catalytically-dependent manner. These results suggest that SMURF2 and USP7 form a regulatory complex that controls site-specific ubiquitination of KAP1, providing dynamic and reversible control over its post-translational modifications and SUMOylation function.

Additionally, our findings expand the role of SMURF2 beyond its canonical function as a degradation-promoting enzyme, establishing it as a key regulator of KAP1 SUMO E3 ligase activity through non-degradative ubiquitination and revealing SMURF2 as a critical modulator of post-translational crosstalk on KAP1. Similar regulatory roles of non-degradative ubiquitination have been reported in other chromatin-associated factors, including histones^51–53^ and, more recently, in the regulation of chromatin occupancy of the NCoR/SMRT/HDAC3 corepressor complex via non-proteolytic ubiquitination of its TAB2 component^54^, highlighting the broader importance of this modification in chromatin regulation.

Having established this regulatory mechanism, we next explored its functional consequences on chromatin and cellular phenotypes. We found that disruption of SMURF2/USP7-mediated regulation of KAP1 substantially alters the epigenetic landscape, leading to global changes in chromatin structure and gene expression. These alterations affect both protein-coding and non-coding genes, including retrotransposons and lncRNAs, underscoring the crucial role of KAP1’s post-translational regulation in maintaining epigenetic homeostasis, particularly in heterochromatin maintenance. The markedly diminished interaction between KAP1-TM and SETDB1/HP1α (Fig. 4L, M) suggests that KAP1-TM either cannot effectively associate with these proteins or forms unstable complexes that fail to support heterochromatin structure. This defect may stem from an impaired ability of KAP1-TM to form homodimeric or oligomeric complexes, which are essential for its interactions with binding partners and its SUMOylation activity^55–57^. However, our biochemical analysis indicates that KAP1-TM retains its ability to form these complexes (Fig. S5), suggesting that the reduced SUMOylation activity and weakened association with chromatin modifiers are more likely due to conformational changes in KAP1-TM, rather than defects in its dimerization.

Additionally, our results demonstrate that altered regulation of KAP1 reshapes its PPI network, which, together with transcriptomic changes, affects several key pathways critical for maintaining cellular function, such as proliferation, invasion, and tumorigenesis. These phenotypes have previously been associated with the expression levels of KAP1^11–13^, highlighting the relevance of our findings. Furthermore, both KAP1 knockdown and TM mutation elicited similar changes in gene expression and cellular phenotypes (Figs. 5A, B; S2D; 7E, F; S4G, H). The most prominent effect was a reduction in cell invasion, observed in both siKAP1 and KAP1-TM cells (Figs. 7E, F; S4G, H), suggesting that this phenotype depends on KAP1’s SUMOylation activity. In line with this, RNA-Seq and RT-qPCR analyses revealed a marked decrease in the expression of MMP7, a key metalloprotease involved in cellular invasiveness, in KAP1-TM cells. Altered expression of other genes and disruptions in KAP1’s interaction network may also contribute to the reduced malignancy of KAP1-TM cells.

Moreover, RNA-Seq and PPI analyses revealed that KAP1-TM disrupts essential cellular programs by altering key pathways involved in cell growth, metabolism, proteostasis, and stress response. Furthermore, comparison of the KAP1-TM and KAP1-WT interactomes, together with RNA-Seq data analysis, indicates that PPI alterations in KAP1-TM cells may represent a compensatory mechanism to buffer, at least in part, transcriptional dysregulation and support altered chromatin functions when KAP1 regulation and its SUMOylation activity are compromised. This adaptation appears to initiate survival-oriented reprogramming, including activation of stress and immune signaling pathways, enhanced RNA processing, and modulation of chromatin-associated processes. Nevertheless, some cellular functions seem to be inadequately compensated, contributing to the altered phenotypes observed in KAP1-TM cells. Collectively, our findings establish the SMURF2/USP7-KAP1-SUMOylation axis as a central regulatory hub integrating ubiquitination and SUMOylation signals to safeguard chromatin architecture and gene expression. The results also reveal the essential role of this axis in regulating the tumorigenic potential of cancer cells. Given its pivotal role in chromatin regulation and tumorigenesis, this axis may represent a novel therapeutic target in cancers characterized by epigenetic dysregulation.

## MATERIALS AND METHODS

### Cell cultures

HEK-293T cells were obtained from the American Type Culture Collection (ATCC). U2OS cells were kindly provided by Prof. Yosef Shiloh (Tel Aviv University, Israel). Cell line authentication was performed by short tandem repeat (STR) profiling at the Genomic Center of the Biomedical Core Facility (Technion, Israel), and cells were routinely tested for mycoplasma contamination. Cells were cultured in high-glucose DMEM (Biological Industries, Israel) supplemented with 2 mM L-glutamine, 10% (v/v) fetal bovine serum (FBS), and 1% (v/v) penicillin-streptomycin. All cultures were maintained at 37°C in a humidified incubator with 5% CO₂.

### Vectors, constructs and site-directed mutagenesis

For transient protein expression, the full-length FLAG-tagged KAP1 (FLAG-KAP1) was generated using the pRK2-FLAG vector backbone, as previously described^34^. Site-directed mutagenesis (SDM) was performed using the Q5 Site-Directed Mutagenesis Kit (NEB, Cat# E0554) to substitute lysines K254, K319 and K779 with arginine (K-to-R), either individually, in double combinations, or as a triple mutant (TM). For stable expression, KAP1-WT and KAP1-TM were subcloned into the pcDNA3-FLAG vector. Primer sequences used for SDM and KAP1 cloning are listed in Table S17. Other vectors and constructs, including MYC-SMURF2 (WT and its E3 ligase-dead mutant), HA-Ubiquitin, and corresponding empty vectors, were previously described^41^. The V5-tagged USP7 and the corresponding empty vector were obtained from VectorBuilder Inc. All constructs were sequence-verified.

### Protein production and purification

GST-SMURF2 (WT and mutant) were produced in *E. coli* and purified using Glutathione Sepharose™ 4B beads (GE Healthcare, Cat# 17075601), as described^58^. FLAG-KAP1 was produced in and affinity-purified from HEK-293T cells, using an approach described in^58,59^, with several modifications. In brief, HEK-293T cells transiently expressing FLAG-KAP1 were lysed using an 0.5%NP-40 lysis buffer (0.5% Nonidet P40 (NP-40) substitute; 25 mM Tris-HCl, pH 7.5; 150 mM NaCl), supplemented with 5 µM ZnSO_4_ and EDTA-free protease inhibitors (Roche, Cat# 11836170001). The samples were then mechanically disrupted by passing through a 29G needle, incubated on ice and centrifuged. The obtained supernatants were incubated with anti-FLAG^®^ M2 affinity gel (Sigma-Aldrich, Cat. #A2220) (4°C, overnight), washed several times with a lysis buffer and resuspended in PBS for subsequent analysis. The purity of the proteins was assessed using PageBlue/Coomassie protein staining (Thermo Fisher Scientific, Cat# 24620).

### Cell transfections and generation of KAP1-depleted and protein-replaced cells

For transient protein expression, HEK-293T and U2OS cells were transfected with the indicated vectors and constructs using polyethylenimine (PEI; Sigma-Aldrich, Cat# 408727) or FuGENE® HD (Promega, Cat# E2312), respectively, according to the manufacturer’s instructions.

To generate stable U2OS cell lines expressing FLAG-KAP1-WT or FLAG-KAP1-TM, cells were first infected with lentiviral particles encoding either an shRNA targeting the 3′UTR of KAP1 (shKAP1) or a control shRNA targeting luciferase (shLuc), using the pLKO.1-KAP1-puro or pLKO.1-Luc-puro vectors, respectively. Infected cells were then selected with puromycin (0.3 μg/ml) for at least two weeks. Following confirmation of stable KAP1 depletion, cells were transfected with either FLAG-KAP1-WT or FLAG-KAP1-TM and selected with G418 (250 μg/ml) for at least one month to establish stable KAP1-reconstituted cell lines.

For transient knockdown of KAP1, predesigned Dicer-substrate siRNA (DsiRNA) duplexes targeting KAP1 or a non-silencing control siRNA (both from Integrated DNA Technologies, IDT) were transfected using Oligofectamine (Invitrogen, Cat# 12252011) according to the manufacturer’s protocol. Cells were analyzed 72 hours post-transfection. The targeting sequences for siNS, siKAP1, shLuc, and shKAP1 are listed in Table S17.

### Cell lysis, western blot analysis, immunoprecipitation, and co-immunoprecipitations

Whole cell lysates (WCL) were prepared by cell lysis in RIPA buffer (50 mM Tris-HCl, pH 7.8; 150 mM NaCl; 1% NP-40 substitute, 0.1% SDS; 0.5% sodium deoxycholate), freshly supplemented with a protease inhibitor cocktail (Roche, Cat# 11836153001) and phosphatase inhibitors (Sigma-Aldrich, Cat# P5726 and P0044), as well as with the deubiquitinase inhibitor N-ethylmaleimide (NEM, 10 mM). Following incubation on ice for 30 min, the samples were sonicated for 1 min (30% amplitude; 4°C) and clarified by centrifugation (14,000 rpm, 15 min, 4°C). The resulting supernatants were collected, and the protein concentration was assessed using QPRO BCA assay kit (Cyanagen, Cat# PRTD1).

For analysis of histone posttranslational modifications, chromatin fractions were extracted using one of the followings methods, as previously described^40^, with some modifications: (i) Cells were collected and lysed in SDS lysis buffer (2% SDS; 10 mM Tris-HCl, pH 7.8; 150 mM NaCl) and immediately boiled at 95°C for 10 min. The lysates were then diluted by adding nine volumes of dilution buffer (50 mM Tris-HCl, pH 7.8; 150 mM NaCl; 1% NP-40 substitute; 0.5% sodium deoxycholate), resulting in a final SDS concentration of 0.2%. The samples were then sonicated (1 min; 30% amplitude; 4°C), clarified by centrifugation, and preserved for subsequent analysis. (ii) Acid based histone extraction. Cell pellets were lysed in hypotonic buffer (10 mM Tris-HCl, pH 8.0; 1 mM KCl; 1.5 mM MgCl₂; 0.2% Triton-X; 1 mM dithiothreitol (DTT); 10 mM sodium butyrate; protease and phosphatase inhibitors) for 30 min at 4°C with gentle mixing. Following the incubation, the nuclei were collected by centrifugation, washed with hypotonic buffer (without Triton-X), and re-suspended in 0.4N H_2_SO_4_. Subsequently, the samples were rotated for 30 min (4°C) and centrifuged (14,000 rpm, 10 min, 4°C). The supernatants were then collected, transferred into fresh tubes, and trichloroacetic acid (TCA) was added dropwise to the histone-containing samples. Following incubation (30 min, on ice), histones were pelleted by centrifugation (14,0000 rpm, 10 min, 4°C) and the supernatants were carefully removed. The resulting pellets were washed twice with ice-cold acetone, air-dried at room temperature for 20 min, and dissolved in nuclease-free ddH_2_O. Protein concentrations in the samples were measured using the BCA assay kit.

For Western blot analysis, equal amounts of proteins from each sample were resolved by SDS-PAGE and transferred for immunoblotting. The list of primary and the corresponding secondary antibodies is provided in Table S17. Immunoblots were detected using WesternBright^™^ ECL reagent (Advansta, Cat# 71-5500) and visualized with the SyngeneG:BOX gel documentation system. Densitometric analysis of immunoblots was performed using Gel.Quant.NET (http://biochemlabsolutions.com/GelQuantNET.html), with signal intensities normalized to appropriate loading controls.

For immunoprecipitations and co-immunoprecipitations, cells were lysed in either RIPA or 1% NP-40 lysis buffer (1% NP-40 substitute; 25 mM Tris-HCl, pH 7.5; 137 mM NaCl; 1 mM EDTA; 1 mM EGTA; 5% glycerol), supplemented with protease and phosphatase inhibitors. Lysates were then sonicated and FLAG-KAP1 was immunoprecipitated using anti-FLAG^®^ affinity gel. Subsequently, beads were washed several times with an ice-cold lysis buffer and boiled for 5 min in 5X SDS sample buffer (50 mM Tris-HCl, pH 8.0; 5 mM EDTA; 5% SDS; 50% glycerol; 50 mM DTT; 0.05% w/v bromophenol blue; 6% β-mercaptoethanol). Immunoprecipitated complexes were resolved by SDS-PAGE and analyzed by immunoblotting with the indicated antibodies.

### Ubiquitination, deubiquitination, and SUMOylation assays

In cellulo ubiquitination assays were performed by co-transfecting HEK-293T cells with FLAG-KAP1 and MYC-SMURF2 (wild-type or mutant), in the presence or absence of HA-ubiquitin. After 24 hours, the cells were harvested and lysed in RIPA buffer supplemented with 10 mM NEM to preserve ubiquitination. The lysates were then incubated on ice for 30 min and sonicated (30% amplitude, 1 min on ice). After clarification by centrifugation, the supernatants were incubated with anti-FLAG^®^ beads to immunoprecipitate FLAG-KAP1. Ubiquitinated KAP1 was detected by immunoblotting using an anti-HA antibody (Thermo Fisher, Cat# 71-5500).

*In vitro* ubiquitination assays were conducted using either purified FLAG-KAP1 (immobilized on agarose beads following affinity purification from HEK-293T cells) or MYC-KAP1 (Origene, Cat# TP301205). These proteins were co-incubated with the components of the ubiquitination cascade: UBE1 (400 ng; Boston Biochem, Cat# E-305), UBE2D3 (600 ng; Boston Biochem, Cat# E2-627), SMURF2 (1 µg GST-SMURF2; WT or E3 ligase-inactive mutant), HA-ubiquitin (20 µg; Boston Biochem, Cat# U-110), and Mg-ATP (4 mM; Boston Biochem, Cat# B-20), in E3 ligase reaction buffer (Boston Biochem, Cat# B-71) for 1.5 hour at 37°C. Beads were then washed four times with ice-cold wash buffer (0.5% NP-40 substitute; 25 mM Tris-HCl, pH 7.5; 150 mM NaCl; 2.5 µM ZnSO_4_; and EDTA-free protease inhibitors) and KAP1 was eluted by boiling in 5X SDS sample buffer.

MYC-KAP1 was pulled down from the reaction using an anti-MYC antibody (Santa Cruz Biotechnology, Cat# sc-40) and protein G Sepharose 4 Fast Flow resin (Cytiva, Cat# 17061801). For KAP1 deubiquitination analysis, we first performed an *in vitro* ubiquitination reconstitution assay, using purified FLAG-KAP1 and the components of the ubiquitination cascade, as described above. The samples were then extensively washed (four times with ice-cold 0.5% NP-40 buffer, and one time with PBS) and resuspended in the deubiquitination buffer (50 mM Tris-HCl, pH 8.0; 50 mM NaCl; 1 mM EDTA; 10 mM DTT; 5% glycerol), supplemented with 5 µM ZnSO_4_. His_6_-USP7 (100 nM; R&D, Cat# E-519) was added to the samples and the reaction was carried out for 1.5 hours at 37°C. Reactions were terminated by adding 0.5% NP-40 buffer (0.5% NP-40 substitute; 25 mM Tris-HCl, pH 7.5; 137 mM NaCl; 1 mM EDTA; 1 mM EGTA; 5% glycerol). Subsequently, the samples were washed five times with ice-cold 0.5% NP-40 buffer and FLAG-KAP1 was then eluted from the beads by boiling in 5X SDS sample buffer. Immunoblotting was performed using the indicated antibodies.

KAP1 auto-SUMOylation assays were performed using purified FLAG-KAP1 proteins (WT and TM), immobilized on beads, and the SUMOylation Assay Kit (Abcam, Cat# ab139470), according to the manufacturer’s instructions. Purified RanGAP1, with or without Mg-ATP, served as a positive or negative control, respectively. Reactions were incubated at 37°C for 2 hours. Subsequently, samples were extensively washed with ice-cold 0.5%NP-40 lysis buffer, and KAP1 was eluted from the beads by boiling in 5X SDS sample buffer. SUMOylation of KAP1 was detected by immunoblotting using the anti-SUMO1 antibody included in the kit. For IRF7 SUMOylation analysis, HEK-293T cells were transfected with either FLAG-KAP1-WT or KAP1-TM. Cells transfected with an empty FLAG vector served as an additional control. Twenty-four hours after transfection, cells were lysed in RIPA buffer, supplemented with protease and phosphatase inhibitors and 10 mM NEM. IRF7 was immunoprecipitated using an anti-IRF7 antibody (Abcam, Cat# ab238137), and its SUMOylation was assessed by Western blotting using anti-SUMO2/3 antibody from the SUMOylation kit.

### Mapping SMURF2-mediated ubiquitination sites on KAP1 using LC-MS/MS

To identify SMURF2-mediated ubiquitination sites on KAP1, we employed three complementary approaches, as described in^58,59^ and outlined in Fig. 1 (panels A, C, E). These approaches included: (i) an *in vitro* ubiquitination assay, (ii) an *in-cellulo* ubiquitination assay without enrichment of KAP1-ubiquitinated proteins, and (iii) an *in-cellulo* assay with enrichment of ubiquitinated KAP1. All assays were performed as detailed in the previous section, followed by MS analysis. The enrichment procedure is described below.

Briefly, FLAG-KAP1 purified from the *in-cellulo* ubiquitination assay was washed with high-stringency buffer (10 mM Tris-HCl, pH 8.0; 1 M NaCl; 1% NP-40 substitute; 1 mM EDTA) and then eluted from FLAG beads using 3X FLAG peptide (450 µg/ml; Sigma-Aldrich, Cat# F4799) in the elution buffer (50 mM HEPES, pH 7.3; 250 mM NaCl; 0.5% NP-40 substitute). The eluates were subjected to filter-aided sample preparation (FASP) for peptide generation prior to LC-MS/MS analysis as follows. Samples were first concentrated and purified using 10 kDa cutoff filter units (Millipore), washed with digestion buffer (50 mM Tris-HCl, pH 8.0), and reduced with DTT. After additional washes, samples were treated with 0.1 M NEM in digestion buffer and digested overnight at 37°C, using sequencing-grade modified trypsin (Promega, Cat# V5111) at a 10:1 protein-to-trypsin ratio (w/w). Following digestion, peptides were washed twice with 0.5 M NaCl and desalted using C18 TopTip cartridges (Glygen, Cat# TT1C18). Samples were then vacuum-dried and reconstituted in immunoaffinity purification (IAP) buffer (50 mM MOPS, pH 7.2; 10 mM sodium phosphate; 50 mM NaCl). Ubiquitinated peptides containing the di-glycine (K-ε-GG) remnant were enriched using the PTMScan® Ubiquitin Remnant Motif (K-ε-GG) kit (Cell Signaling, Cat# 5562) by overnight incubation at 4°C (with rotation). Subsequently, beads were washed twice with 0.25% NP-40 substitute in IAP buffer, twice with IAP buffer alone, and twice with MS-grade water. The bound peptides were then eluted with 0.15% trifluoroacetic acid under acidic conditions and vortexed at low speed for 10 min.

Finally, peptides were desalted using NuTip C18 pipette tips (Glygen, Cat# NT2C18), lyophilized and submitted for LC-MS/MS analysis at the Smoler Proteomics Center (Technion, Israel). The MS data were analyzed using MaxQuant (https://www.maxquant.org/).

### Stratification of the KAP1-WT and KAP1-TM interactomes by LC-MS/MS and data analysis

The analysis was conducted using a previously established workflow^34,58,59^, with some modifications. Briefly, HEK-293T cells were transfected with the indicated constructs and lysed in 1% NP-40 lysis buffer supplemented with protease and phosphatase inhibitors, as described above. The lysates were then precleared by incubation with protein A/G PLUS-agarose beads (Santa Cruz, Cat# sc-2003) for 2 hours at 4°C with rotation, and FLAG-KAP1 was immunoprecipitated using anti-FLAG^®^ M2 affinity gel (4°C, overnight). Subsequently, the beads were washed four times with ice-cold lysis buffer, twice with pre-urea wash buffer (50 mM Tris, pH 8.5; 1 mM EGTA; 75 mM KCl), and four times with PBS. The elution buffer (8 M urea; 20 mM Tris, pH 7.5; 100 mM NaCl) was then added to the beads and the samples were incubated on the shaker for 30 min (room temperature). The beads were then pelleted and the supernatants, containing the eluted proteins, were transferred to fresh tubes. The samples were sent to the Smoler Proteomics Center (Technion, Israel), where they were trypsin-digested and analyzed using LC-MS/MS in Q Exactive^™^ Plus mass spectrometer (Thermo Fisher Scientific). The peptide identification and annotation were carried out using the Proteome Discoverer™ software (Thermo Fisher Scientific), with MS Amanda and Sequest^™^ HT search engines. All identified peptides were filtered for high confidence: FDR < 1%, top rank, high mass accuracy, and a minimum coverage of two peptides per protein. The resulting list of KAP1 interactors was subjected to GO analysis using the DAVID functional annotation tools (https://davidbioinformatics.nih.gov/), accessed on February 10, 2024. GO terms with Benjamini-Hochberg-adjusted q-values < 0.05 were considered as statistically significant.

Interactors were classified into functional categories using the PANTHER 18.0 tool (https://pantherdb.org/), which was also used for pathways analysis (PANTHER Pathway v3.6.7). The protein-protein interaction networks were constructed using the STRING v11.5 tool (https://string-db.org/), based on high-confidence evidence from curated databases and experimental data, as well as predicted interactions (e.g., gene neighborhood, fusions, co-occurrence) and additional sources (e.g., text mining, co-expression and protein homology). Only interactors with established network connections were included in the final analysis. *k*-means clustering was applied to classify proteins into distinct functional groups.

### KAP1 dimerization/oligomerization assay

HEK-293T cells, expressing either FLAG-KAP1-WT or KAP1-TM, were lysed in a non-reducing, non-denaturing buffer (50 mM Tris-HCl, pH 7.5; 150 mM NaCl; 1 mM EDTA; 1% NP-40 substitute; 0.1% Triton-X), supplemented with protease and phosphatase inhibitors. The resulting cell extracts were mixed with 5X native sample buffer (50 mM Tris-HCl, pH 8.0; 5 mM EDTA; 27% glycerol; 0.05% bromophenol blue), diluted to a final 1X concentration, and resolved on a 6% native PAGE gel. The gel electrophoresis was performed using native running buffer (25 mM Tris-HCl; 192 mM glycine; pH 8.3), and KAP1 dimerization/oligomerization was analyzed by immunoblotting with anti-KAP1 antibody (Abcam, Cat# ab109545). In parallel, aliquots of the same lysates were mixed with 5X SDS sample buffer (diluted to a final 1X concentration), boiled and resolved in SDS-PAGE to detect the monomeric forms of the proteins, as described above.

### Cell proliferation, migration, and invasion assays

These assays were conducted as previously described^40,41^ with minor modifications, as detailed below. To assess short-term cell proliferation, KAP1-modified cells were seeded at equal density (10,000 cells per 60 mm plate) and cultured for the indicated time periods. At each time point, cells and their supernatants were collected, centrifuged, stained with 0.2% trypan blue solution (Biological Industries, Cat# 03-102-1B), and analyzed using the Cellometer^™^ Auto T4 Cell Counter (Nexcelom Bioscience).

To evaluate long-term proliferative capacity and colony formation potential, cells were seeded at equal density (4,000 cells per 60 mm plate) and cultured for 12 days, with medium replaced every 4 days. At the end of the incubation period, cell colonies were stained with 2% crystal violet solution (prepared in 50% ethanol; MP Biomedicals, Cat# 0210177525), visualized under a light microscope, photographed, and counted to determine both the number of transformed colonies and the total number of colonies.

To assess cell migration, cells were seeded at equal density (50,000 cells) into the upper chamber of 8.0-µm pore transwell inserts (Greiner Bio-One, Cat# 662638) in serum-free DMEM and incubated at 37°C for 24 hours. The lower chamber was filled with complete growth medium. After incubation, non-migrated cells on the upper surface of the membrane were gently removed using a cotton swab. Membranes were then fixed with 3.7% formaldehyde for 2 min, followed by PBS washes. Subsequently, membranes were permeabilized with 100% methanol for 20 min, washed with PBS, and stained with 0.5% crystal violet solution for 15 min. After additional PBS washes, the membranes were air-dried overnight.

For the invasion assay, cells were seeded in serum-free DMEM into the upper chamber of 8.0-µm pore transwell inserts pre-coated with growth factor-reduced Matrigel basement membrane matrix (2.5 mg/ml; Corning, Cat# 354234). The lower chamber was filled with complete growth medium. After 24 hours, membranes were processed as described for the migration assay. In both assays, the stained membranes were imaged using an Eclipse TS100 inverted phase contrast microscope (10x/0.25 NA air objective; Nikon). The crystal violet stain was then eluted with 1% SDS and quantified using a spectrophotometer at 570 nm, as previously described^40^.

### Cell cycle analysis by bi-parametric flow cytometry

This analysis was conducted as described^40,41^, with several adjustments. Briefly, exponentially growing cells were pulsed with 20 µM bromodeoxyuridine (BrdU; Sigma-Aldrich, Cat# B5002) for 1 hour at 37°C, then harvested, washed with ice-cold PBS and pelleted by centrifugation (1,200 rpm, 5 min, 4°C). Cells were resuspended in 1 ml PBS, fixed by dropwise addition of 4 ml cold (-20°C) absolute ethanol with gentle vortexing, and then stored at -20°C.

For analysis, fixed cells were washed twice with ice-cold PBS, permeabilized in freshly prepared 2 M HCl containing 0.5% Triton X-100 for 30 min (room temperature), and neutralized in 0.1 M sodium borate buffer (pH 8.5) for 5 min. Cells were then washed with 1% BSA in PBS and incubated with a mouse anti-BrdU monoclonal antibody (Chemicon, Cat# MAB3222; 1:100) in 100 µl PBS containing 1% BSA, 0.25% Tween-20, and 2 µg RNase A (Sigma-Aldrich, Cat# R6513) for 30 min (at room temperature). Following two washes in 1% BSA/PBS, cells were incubated with Alexa Fluor 488-conjugated anti-mouse IgG (Jackson ImmunoResearch, Cat# 715-545-150; 1:200) in 1% BSA/0.25% Tween-20/PBS for 25 min at room temperature, protected from light. After two additional washes, cells were resuspended in 500 µl PBS and transferred to 12 x 75 mm polystyrene tubes with mesh caps. Subsequently, propidium iodide was added to a final concentration of 10 µg/ml, and cells were incubated at room temperature, protected from light. BrdU incorporation and DNA content were assessed by bi-parametric flow cytometry using a Gallios flow cytometer (Beckman Coulter). Data were analyzed using the Gallios software.

### Immunofluorescence and confocal microscopy

Cells growing on poly-D-lysine-coated glass coverslips were fixed with 4% formaldehyde for 20 min at room temperature. After fixation, cells were washed with PBS containing 0.1% Tween-20 (PBST) and permeabilized with 0.5% Triton X-100 for 20 min at room temperature. To block non-specific binding, cells were incubated in blocking solution consisting of 3% BSA in 0.1% Tween-20/PBS for 1 hour. Subsequently, cells were incubated with primary monoclonal antibodies: mouse anti-FLAG (Sigma-Aldrich, Cat# F3165; 1:1000) and rabbit anti-histone H3 (trimethyl K9) antibody (ChIP grade; Abcam, Cat# ab176916; 1:1000) for 2 hours at room temperature. Following washes with PBST, cells were incubated in the dark for 1 hour with fluorophore-conjugated secondary antibodies: Alexa Fluor^®^ 488 donkey anti-rabbit IgG (Jackson ImmunoResearch, Cat# 711-546-152; 1:1000) and Rhodamine Red™-X goat anti-mouse IgG (Jackson ImmunoResearch, Cat# 115-296-071; 1:500). After additional washes, cell nuclei were counterstained with Hoechst 33258 (1 µg/ml, 5 min; Sigma-Aldrich, Cat# B2883). Coverslips were then mounted onto glass slides using ProLong™ Diamond Antifade Mountant (Thermo Fisher Scientific, Cat# P36961) and imaged using an LSM780 inverted confocal microscope (Zeiss) with a Plan-Apochromat 63×/1.40 Oil DIC M27 objective. All comparative images were acquired under identical microscope and camera settings. Image analysis was performed using Zeiss ZEN Blue (v2.3 lite) and ImageJ software tools (https://imagej.net/ij/).

### Micrococcal nuclease (MNase) sensitivity assay

This assay was performed as previously described^40,41^, with some modifications. Briefly, nuclei were isolated from KAP1-replaced cells using a hypotonic lysis buffer (10 mM HEPES, pH 7.9; 10 mM KCl; 1.5 mM MgCl_2_; 0.34 M sucrose; 10% glycerol; 0.1% Triton X-100; 1 mM DTT) supplemented with freshly added EDTA-free protease inhibitor cocktail (Roche, Cat# 11836170001) and phosphatase inhibitors (Sigma-Aldrich, Cat# P5726 and P0044). Cells were lysed at 4°C for 5 min and then centrifuged. The nuclear pellet was washed once with hypotonic buffer without Triton X-100, and nuclear yield was determined by resuspending an aliquot in 1N NaOH and measuring absorbance at 260/280 nm. Equal amounts of nuclei (100 μg) were resuspended in MNase digestion buffer (15 mM Tris-HCl, pH 7.4; 60 mM KCl; 0.25 M sucrose; 15 mM NaCl; 1 mM CaCl_2_; 0.5 mM DTT) and digested with micrococcal nuclease S7 (Roche, Cat# 10107921001; 3 U per 16 µg DNA) at room temperature for the indicated times. Nuclei incubated in digestion buffer without MNase served as controls. Reactions were terminated by adding an equal volume of 2X TNESK buffer (20 mM Tris-HCl, pH 7.4; 0.2 mM NaCl; 2 mM EDTA; 2% SDS) freshly supplemented with 0.2 mg/ml proteinase K (Qiagen, Cat# 158918), followed by overnight incubation at 37°C. DNA was purified by phenol-chloroform extraction, resuspended in TE buffer (10 mM Tris-HCl, pH 8.0; 1 mM EDTA), and resolved on a 1.5% agarose gel stained with SYBR Safe (Invitrogen, Cat# S33102). Gel images were quantified using GelQuant.NET (http://biochemlabsolutions.com/GelQuantNET.html).

### RNA sequencing

Total RNA was extracted from KAP1-WT and KAP1-TM U2OS cells using the RNeasy mini kit (Qiagen, Cat# 74004), according to the manufacturer’s instructions. RNA integrity was assessed with the Agilent RNA Pico 6000 Kit and Bioanalyzer 2100 (Agilent Technologies). For each sample, 1 µg of total RNA was used for mRNA enrichment using the NEBNext Poly(A) mRNA Magnetic Isolation Module (NEB, Cat# E7490). RNA-Seq libraries were generated using the NEBNext Ultra II RNA Library Prep Kit for Illumina (NEB, Cat# E7770).

Library concentrations were measured with the Qubit dsDNA HS Assay Kit (Invitrogen/Thermo Fisher Scientific) on a Qubit 2.0 fluorometer, and library quality was verified using the Agilent D1000 ScreenTape System and TapeStation 4200. Subsequently, 250 nM of each library were pooled together and diluted to 4 nM according to the NextSeq system protocol. A final concentration of 1.5 pM, including 1% PhiX control, was loaded onto the flow cell. Sequencing was performed on an Illumina NextSeq 550 platform using single-end 75-cycle reads, following the manufacturer’s protocol. Library preparation and sequencing were conducted at the Faculty Genome Technology Center (Bar-Ilan University, Israel).

### Bioinformatic analysis of RNA-Seq data

Raw FASTQ files generated using the Illumina NextSeq 550 platform were obtained from three independent biological replicates per condition. Reads were aligned to the human reference genome (hg38; GENCODE v32 annotation, September 2019 release) using Bowtie2 within the Partek Flow software (https://www.partek.com). Gene-level read counts were quantified based on the GENCODE v32 annotation, and differential gene expression analysis was performed using DESeq2. Genes showing changes in expression greater than 25%, a false discovery rate (FDR) < 1%, and *P*<0.05 were defined as differentially expressed genes (DEGs). The functional enrichment analysis of DEGs was conducted using the DAVID tool (accessed January 5, 2025). GO terms with *P*<0.05 were considered as significantly enriched.

For the analysis of LINE-1 (L1) expression, raw sequence reads were aligned to the human reference genome (hg38) using the STAR aligner (v2.7.11b). To focus on uniquely mapping reads, sequences aligning to multiple loci were excluded by -outFilterMultimapNmax 1. In cases of multiple alignments, only the read with the highest alignment score was retained using -outFilterMultimapScoreRange 1. A mismatch threshold of 3% per base was applied with -outFilterMismatchNoverLmax 0.03 to ensure alignment quality.

L1 elements were annotated using an L1-specific GTF file downloaded from the UCSC Table Browser data retrieval tool (http://genome.ucsc.edu). Quantification of individual LINE-1 elements was performed using the featureCounts from the Rsubread package (Bioconductor; https://bioconductor.org/packages/release/bioc/html/Rsubread.html). Count normalization was carried out using DESeq2, scaling mapped reads to the reference genome in a replicate-wise manner (https://www.bioconductor.org/packages/release/bioc/html/DESeq2.html).

Classification of L1 elements based on evolutionary age (young, intermediate, and old) was performed based on published criteria^47,48^. Box plots of LINE-1 expression across age groups were generated using the Plotly Python package (https://plot.ly). Heatmaps of individual L1 elements were created using Morpheus (https://software.broadinstitute.org/morpheus/) via the macOS terminal interface (Sonoma 14.6.1).

### KAP1 Chip-Seq data analysis

Raw sequence reads (SRR501957 [input], SRR502577 [replicate 1], SRR502578 [replicate 2])^42^ were processed for adapter trimming using Trimmomatic (http://www.usadellab.org/cms/index.php?page=trimmomatic), and read quality was assessed with FastQC (https://www.bioinformatics.babraham.ac.uk/projects/fastqc/). High-quality reads were aligned to the human reference genome (hg38) using Bowtie2 (http://bowtie-bio.sourceforge.net/bowtie2/index.shtml) with the “very-sensitive-local” alignment parameter. PCR duplicates were removed using SAMtools RmDup (http://samtools.sourceforge.net), and ChIP versus input BAM files were compared using deepTools2 (https://deeptools.ie-freiburg.mpg.de) for matrix computation and data visualization. ChIP-seq peaks were called using MACS2 (http://liulab.dfci.harvard.edu/MACS/) with default parameters. The identified peaks were annotated with HOMER software suite (http://homer.ucsd.edu/homer/) using the human reference genome annotation file obtained from the UCSC genome browser.

### Quantitative real time PCR (RT-qPCR)

RT-qPCR analysis was performed as previously described^60^, using RNeasy mini kit (Qiagen, Cat# 74004). cDNA was synthesized from total RNA using random primers and the High-Capacity cDNA Reverse Transcription Kit (Applied Biosystems, Cat# 4368814). Equal amounts of cDNA (100 ng per reaction) were amplified using Fast SYBR^™^ Green Master Mix (Applied Biosystems, Cat# 4385612) on a ViiA^™^ 7 Real-Time PCR System (Thermo Fisher Scientific). Gene expression was assessed using the 2^−ΔΔCt^ method and normalized to the expression of the housekeeping gene GAPDH (glyceraldehyde 3-phosphate dehydrogenase).

### Animals and xenograft tumor formation assay

NSG mice (NOD.CB17-*Prkdc^Scid^lL2rg^tm1^*/BcgenHsd; Envigo) were housed in a specific pathogen-free (SPF) facility at the Faculty of Medicine, Bar-Ilan University (Israel), in accordance with Federation of Laboratory Animal Science Associations (FELASA) guidelines. To establish the xenograft tumor formation model, 6-8-week-old NSG mice were subcutaneously injected with 2x10^6^ KAP1-WT or KAP1-TM U2OS cells suspended in 200 μl of PBS/Matrigel solution. Cell number and viability before and after injection were assessed using the Cellometer^™^ Auto T4 Cell Counter with trypan blue exclusion. Tumor growth was measured using a vernier caliper, and tumor volume was calculated using the formula: length x width^2^ x 0.5. All animal procedures were approved by the Bar-Ilan University Animal Care and Use Committee (protocol number BIU-MD-IL-2301-105-5).

### Statistical analysis

Data are presented as mean ± standard error of the mean (SEM), unless otherwise indicated. Statistical significance was assessed using unpaired Student’s *t*-test or ANOVA, as appropriate, using Microsoft Excel and IBM SPSS Statistics (v28). *P*-values less than 0.05 were considered statistically significant. Sample sizes, number of replicates, and specific statistical parameters are indicated in the figure legends.

### Data and resource availability

RNA-Seq data (FASTQ files) are available in the NCBI Sequence Read Archive (SRA) under the BioProject accession no. PRJNA1299141. Reviewers can access the dataset through the link: https://dataview.ncbi.nlm.nih.gov/object/PRJNA1299141?reviewer=t6i6886fj5thh8p4kde3q82gvn The mass spectrometry proteomics data for the KAP1-WT and KAP1-TM interactomes have been deposited in the ProteomeXchange Consortium via the PRIDE partner repository, with the dataset identifier PXD067081. Reviewers can access the dataset by logging into the PRIDE website using the username: and the password: tCUCX0kxLKep. The requests for resources can be directed to the lead contact, Dr. Michael Blank.

## ACKNOWLEDGMENTS

We thank members of the Blank lab for valuable discussions and assistance with experiments. We also acknowledge the staff of the Faculty Instrumental Core Facility and the Genome Technology Center for their support with confocal microscopy, flow cytometry, and RNA-Seq analysis. M.B. acknowledges support from the Israel Science Foundation (422/20 and 1460/24) and the Dayan Family Foundation.

## AUTHOR CONTRIBUTIONS

S.B.S., K.N., P.A.S., and G.L.C. performed experiments and analyzed the data. M.S. conducted bioinformatic analysis of RNA-Seq data to identify differentially expressed genes in KAP1-TM versus WT cells, as well as in siKAP1 versus siNT U2OS cells, using a publicly available RNA-Seq dataset^43^. N.R.C. performed bioinformatic analyses of LINE-1 expression using both our RNA-Seq data and publicly available datasets^43^, and also analyzed KAP1 ChIP-Seq data from U2OS cells^42^. M.B. conceived the study, designed the research strategy, supervised the project, and wrote the manuscript.

