## Supplemental Figures S1-S5, Supplemental Tables S1-S2, and Supplemental Figure and Table Legends for "SMURF2/USP7-mediated ubiquitination of KAP1 controls its SUMO E3 ligase activity and chromatin regulation"

**Supplemental Figure Legends:** pages 2-5

**Supplemental Figures S1–S5:** pages 6-10

**Supplemental Table Legends:** page 11

**Supplemental Tables S1–S2:** pages 12-13

**Supplemental Tables S3–S17:** provided as individual Excel files with the submission

### SUPPLEMENTAL FIGURE LEGENDS

**Figure S1. Generation of KAP1-replaced cells**

**(A)** Schematic diagram of the workflow used for generating KAP1-replaced cells.

**(B)** Western blot analysis showing KAP1 expression in KAP1-replaced U2OS cells. shLuc: shLuciferase control.

**Figure S2. Effects of KAP1 on gene expression and LINE-1 retroelement activity**

**(A)** lncRNAs exhibiting significant differences in expression between KAP1-TM and WT cells. Fold changes for each lncRNA (upregulated or downregulated) are shown on the X-axis. The annotation for each lncRNA was obtained using the LNCipedia (https://lncipedia.org) and GeneCards (https://www.genecards.org) databases.

**(B)** ChIP-Seq analysis of KAP1 chromatin binding in U2OS cells, showing a heatmap of KAP1 occupancy around the transcription start sites (TSS) and within gene bodies. The X-axis represents the distance from the TSS in kilobases (Kb), ranging from -5.0 Kb to +5.0 Kb. The color gradient indicates KAP1 binding density, with the color bar representing the log2-fold enrichment of KAP1 relative to the input control.

**(C)** The number of genes that were upregulated or downregulated in KAP1-TM versus WT cells, containing KAP1 binding sites (in the promoter, gene body, or both). The pie chart (bottom panel) shows the percentage of these genes out of a total of 298 differentially expressed genes in KAP1-TM vs. WT cells.

**(D)** RT-qPCR analysis of selected genes in KAP1-TM versus WT cells. Corresponding gene expression changes from RNA-Seq analyses (KAP1-TM vs. WT and siKAP1 vs. siNT control) are summarized in the table below, along with the presence of KAP1 binding sites at these loci. Data are presented as mean $\pm$ SEM from three biological replicates, each with three technical replicates. *$P < 0.05$; **$P < 0.01$; ***$P < 0.001$. CHRDL1, chordin like 1; MMP7, matrix metalloproteinase 7; MYOCD, myocardin; CREB3L1, cAMP responsive element binding protein 3 like 1; CCBE1, collagen and calcium binding EGF domain containing protein 1.

**(E)** Box plots showing the effect of KAP1 knockdown on the expression of LINE-1 retroelements. Data are mean ± SD (N=3). *$P < 0.05$; **$P < 0.01$. ns, not significant.

**(F)** Heatmaps of the individual LINE-1 elements expressed in KAP1 knockdown vs. control cells.

**Figure S3. KAP1 interactome analysis: complementary results and analysis of KAP1 interactions within the shared protein group**

**(A)** Validation of the efficiency of KAP1 pull-down from KAP1-WT and KAP1-TM-expressing cells for interactome analysis. The right panel shows Western blot analysis of the immunoprecipitated FLAG-KAP1, probed with anti-FLAG and anti-KAP1 antibodies. The left panel displays Coomassie gel staining of KAP1-WT and TM proteins in SDS-PAGE. IP: immunoprecipitation; WCL: whole cell lysates.

**(B)** Functional protein association networks of KAP1-WT and KAP1-TM unique interactors. The PPI maps were generated using the STRING 11.5 platform (https://string-db.org). Clusters identified by *k*-means clustering are shown in different colors. Nodes represent different proteins, with the indicated gene symbols.

**(C)** Venn diagrams showing the total number of KAP1-associated proteins identified in the KAP1-WT and KAP1-TM interactomes. The bottom panel displays KAP1 interactions within the shared protein group after applying a two-fold change threshold in protein abundance.

**(D)** Protein classes of shared KAP1 interactors enriched at ≥2-fold, determined using the PANTHER classification system. The top ten categories with the highest number of interactors are shown. Full data are provided in Supplemental Table S13. Protein classes highlighted in bold indicate a >50% change in abundance between KAP1-WT and KAP1-TM samples.

**(E)** Functional protein association networks of KAP1-WT and KAP1-TM common interactors, constructed using the STRING 11.5 platform.

**(F-H)** GO enrichment analysis of molecular functions, biological processes and cellular pathways in KAP1-TM versus WT cells, based on shared protein interactors enriched more than two-fold. The number of KAP1-associated proteins in each GO category from the KAP1-WT and KAP1-TM interactomes is shown on the X-axis. The top ten categories with the most significant *p*-values are shown for each GO term. Full datasets are provided in Supplemental Tables S14-S16. A total of 39 molecular functions, 74 biological processes and 230 cellular pathways were significantly enriched in the KAP1-WT interactome, compared to only 8 molecular functions, 9 biological processes, and 11 pathways in the KAP1-TM interactome. Note that although some GO terms were shared between the two interactomes, the number of KAP1-associated proteins was significantly reduced in the KAP1-TM interactome.

**Figure S4. Impact of KAP1 knockdown and re-expression on cell proliferation, migration and invasion**

**(A)** Growth curves of U2OS cells stably expressing KAP1-WT and KAP1-TM over a 96-hour period. Data are mean $\pm$ SEM. ns, not significant.

**(B)** Bi-parametric flow cytometry analysis of cell cycle distribution in U2OS cells stably expressing either KAP1-WT or KAP1-TM. Cells were pulsed with 20 µM BrdU for 1 hour and then stained with anti-BrDU antibody and propidium iodide (PI). The upper table summarizes data from three experiments, presented as mean ± SD, with the indicated *p*-values. The lower panel shows representative 2-D flow cytometry plots of KAP1-WT and KAP1-TM cells co-stained with PI and anti-BrdU.

**(C)** Western blot analysis of KAP1 expression in U2OS cells following KAP1 knockdown (siKAP1) and transient re-expression (siKAP1+KAP1-WT and siKAP1+KAP1-TM). siNT:

non-targeting siRNA. siKAP1+vector: KAP1-knockdown cells transfected with an empty FLAG vector.

**(D)** Growth curves of U2OS cells with KAP1 knockdown and transiently re-expressed KAP1. Data are mean $\pm$ SEM.

**(E, F)** Transwell cell migration assays of U2OS cells following transient KAP1 knockdown and/or protein re-expression. Scale bars, 100 μm. Data are mean $\pm$ SEM. ns, not significant.

**(G, H)** Transwell cell invasion assays of U2OS cells with transient KAP1 knockdown and/or protein re-expression. Scale bars, 100 μm. Data are mean $\pm$ SEM. *$P < 0.05$; **$P < 0.01$.

**Figure S5. KAP1-TM retains its ability to oligomerize**

HEK-293T cells were transfected with either KAP1-WT or KAP1-TM. After 24 hours, cells were lysed under native conditions and analyzed by native PAGE (left panel) or by SDS-PAGE following sample denaturation (right panel). Immunoblotting with anti-KAP1 antibody reveales oligomeric KAP1 species (>250 kDa) under native conditions and monomeric forms (~110–120 kDa) under denaturing conditions.

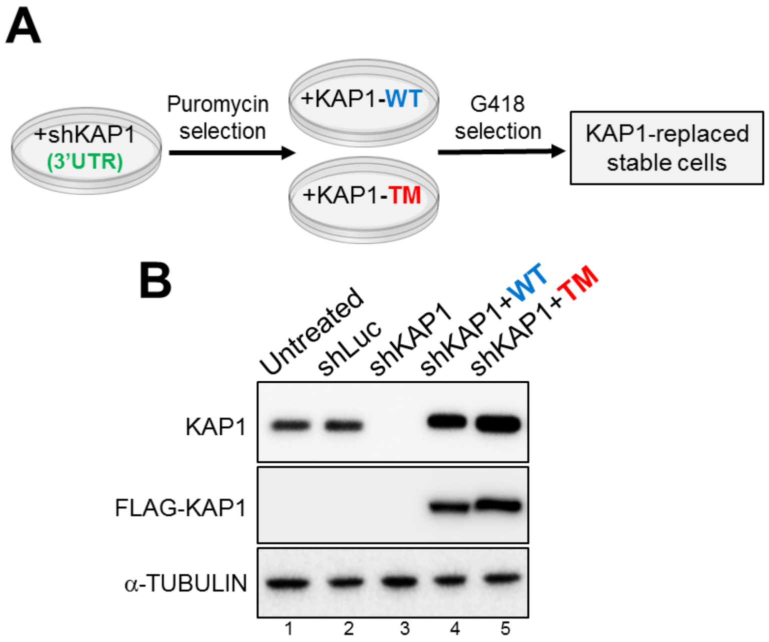


**SUPPLEMENTAL FIG. S1**

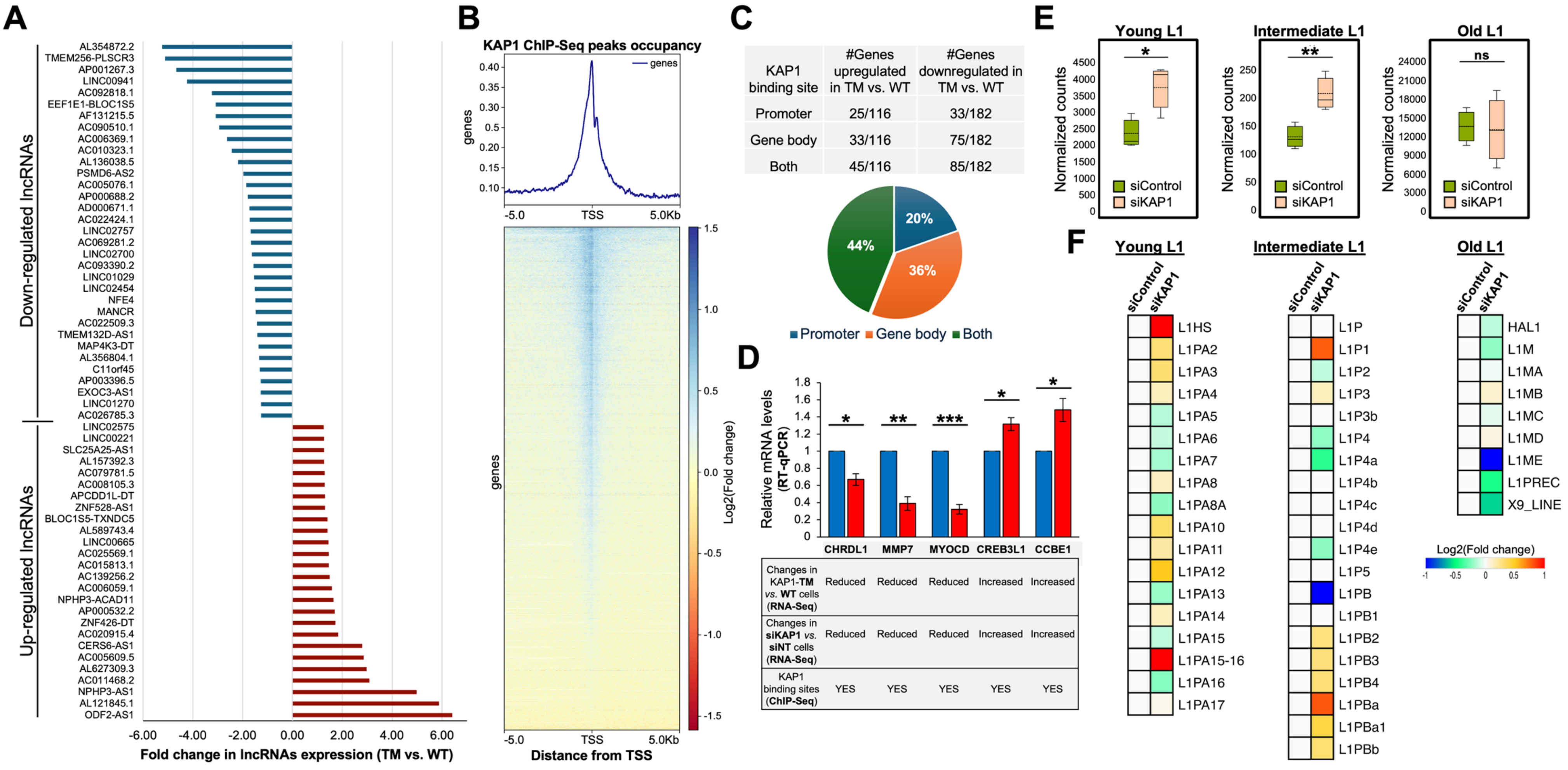


SUPPLEMENTAL FIG. S2

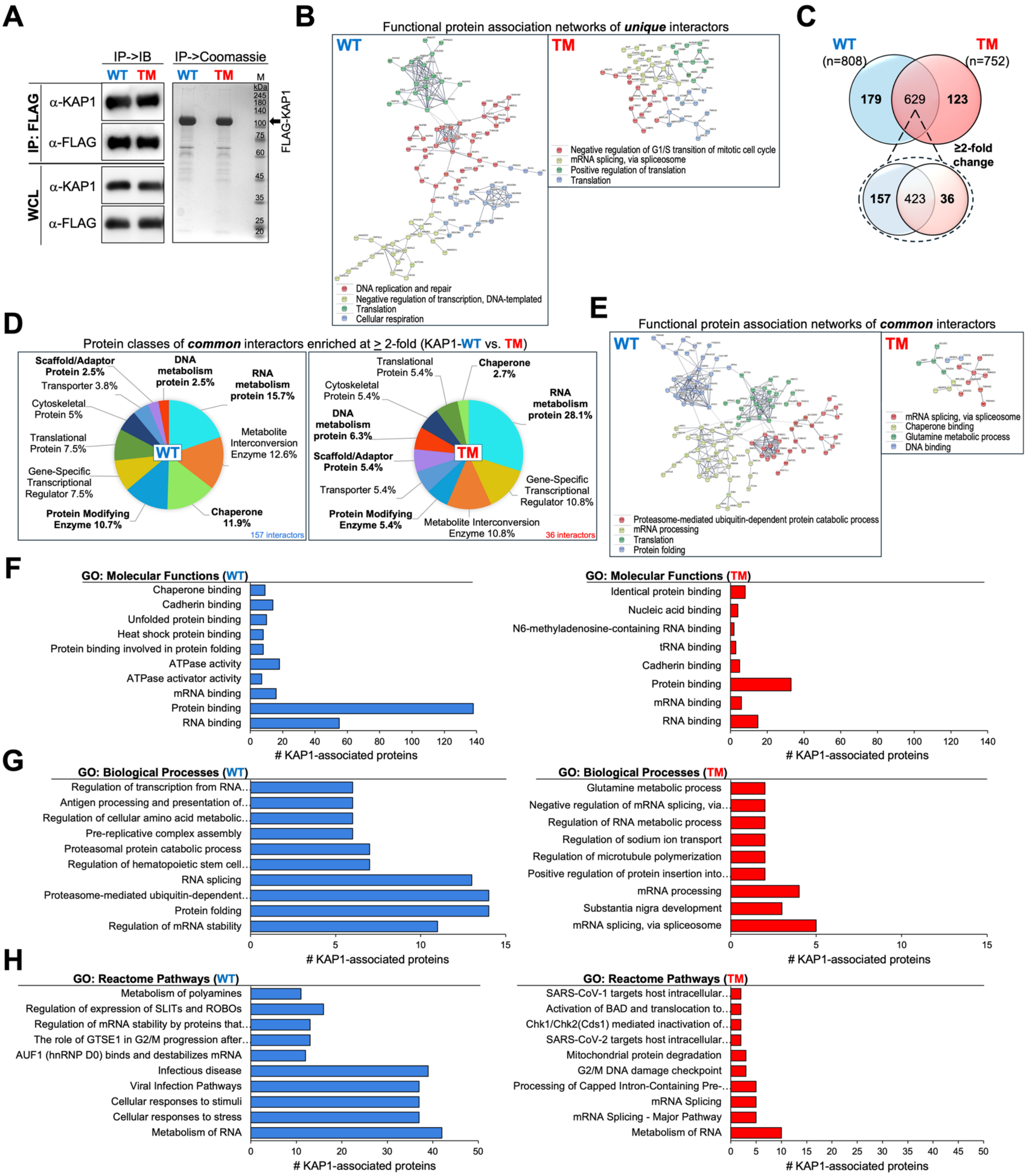


SUPPLEMENTAL FIG. S3

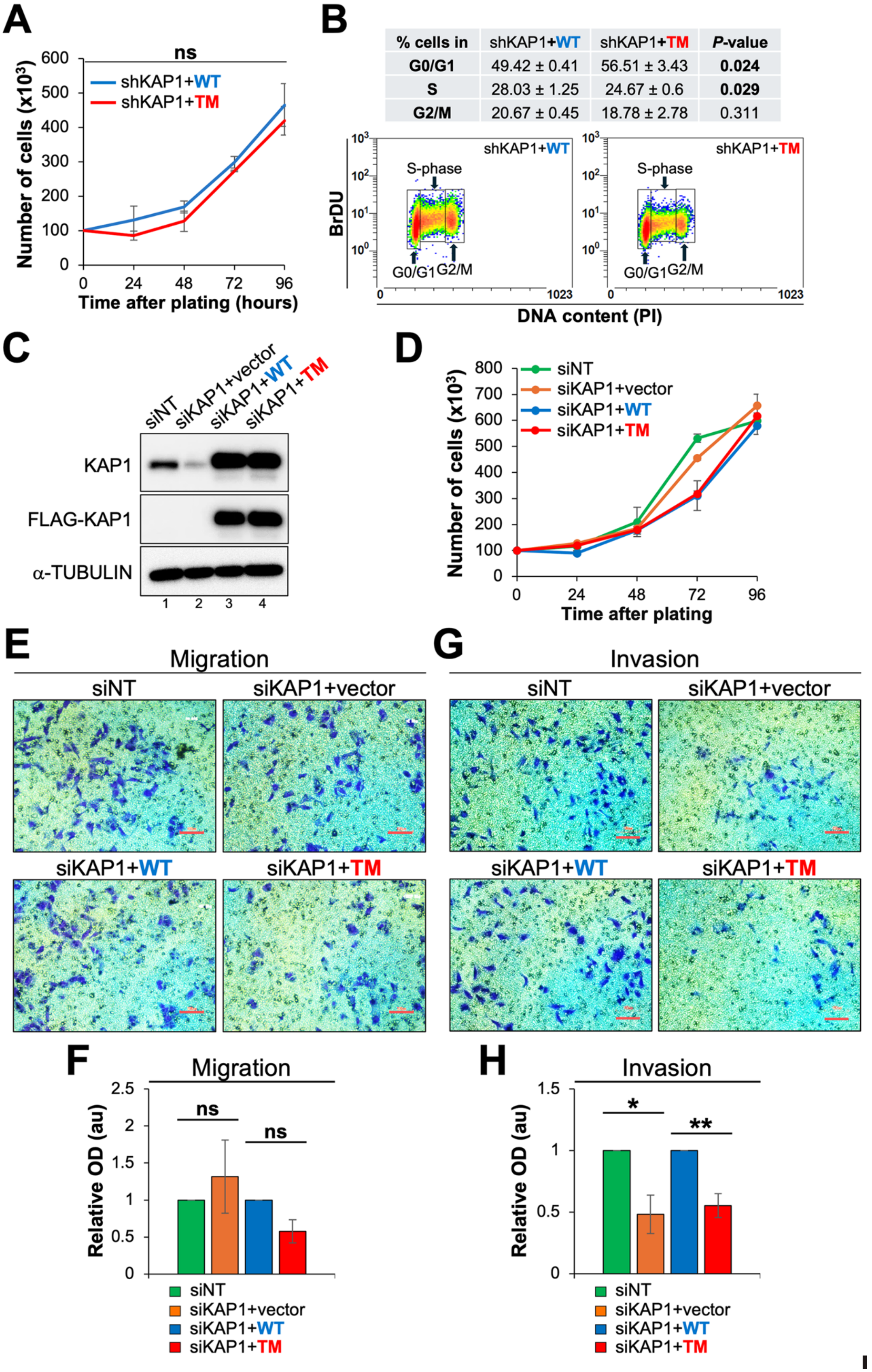


SUPPLEMENTAL FIG. S4

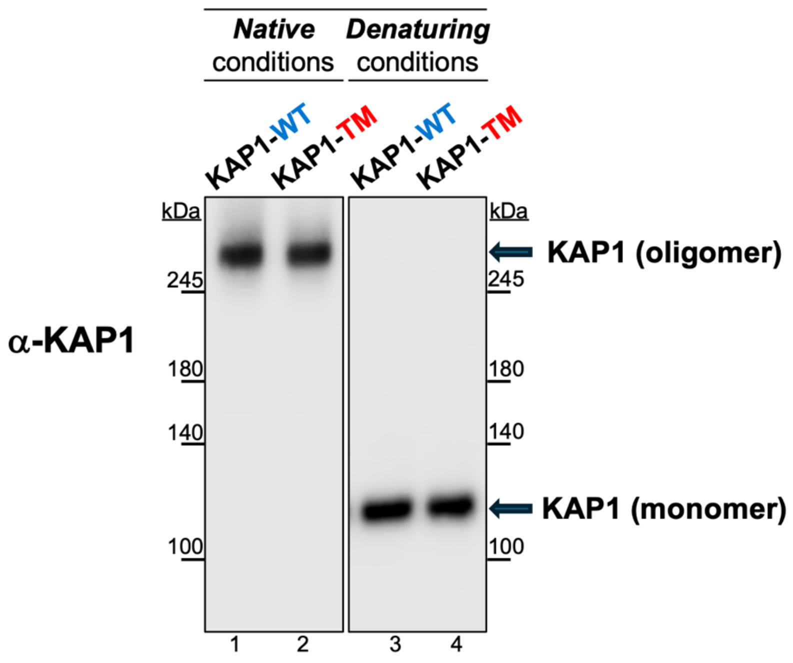


**SUPPLEMENTAL FIG. S5**

### SUPPLEMENTAL TABLE LEGENDS

**Table S1.** SMURF2-mediated ubiquitination sites identified on KAP1 using LC-MS/MS.

**Table S2.** K254, K319 and K779 residues of KAP1 identified in high-throughput proteomic studies as potential ubiquitination sites.

**Table S3.** Genes significantly up- and down-regulated in KAP1-TM vs. WT U2OS cells (RNA-Seq data with applied threshold).

**Table S4.** Differentially expressed genes in KAP1-TM vs. WT cells with annotated KAP1 binding sites.

**Table S5.** Genes differentially expressed in KAP1-TM vs. WT cells with consistent directional changes in KAP1 knockdown (siKAP1) vs. control (siNT) cells**.**

**Table S6.** GO: Molecular functions enriched in either KAP1-TM vs. WT cells (RNA-Seq).

**Table S7.** GO: Biological processes enriched in either KAP1-TM vs. WT cells (RNA-Seq).

**Table S8.** GO: Cellular pathways enriched in KAP1-TM vs. WT cells (RNA-Seq).

**Table S9.** Protein classes of unique interactors for KAP1-WT and KAP1-TM.

**Table S10.** GO: Molecular functions of unique KAP1 interactors in KAP1-TM vs. WT cells.

**Table S11.** GO: Biological processes of unique KAP1 interactors in KAP1-TM vs. WT cells.

**Table S12.** GO: Cellular pathways of unique KAP1 interactors in KAP1-TM vs. WT cells.

**Table S13.** Protein classes of common KAP1 interactors (after two-fold enrichment).

**Table S14.** GO: Molecular functions of common KAP1 interactors (after two-fold enrichment).

**Table S15.** GO: Biological processes of common KAP1 interactors (after two-fold enrichment).

**Table S16.** GO: Cellular pathways of common KAP1 interactors (after two-fold enrichment).

**Table S17.** Key resources table.

**Table S1.** SMURF2-mediated ubiquitination sites identified on KAP1 using MS analysis.

| Peptide sequence with ubiquitinated lysine residue | ABUNDANCE (PEAK AREA) KAP1+ Empty Vector | KAP1+ SMURF2-WT | KAP1+ SMURF2-Mut | Identification method (approach) |
|---|---|---|---|---|
| LTEDkADVQSIIGLQR (**KAP1_K779**) | 0 | **6.41x10$^{7}$** | 0 | In vitro ubiquitination assay and in-gel digestion (approach #1) |
| kLLASLVK (**KAP1_K254**) | 0 | **3.23x10$^{6}$** | 0 | In cellulo ubiquitination assay and in-gel digestion (approach #2) |
| VLVNDAQkVTEGQQER (**KAP1_K319**) | 0 | **2.53x10$^{7}$** | 1.37x10$^{7}$ | In cellulo ubiquitination assay and in-gel digestion (approach #2) |
| VLVNDAQkVTEGQQER (**KAP1_K319**) | 0 | **2.97x10$^{7}$** | 0 | In cellulo ubiquitination assay and K-e-GG enrichment (approach #3) |
| LTEDkADVQSIIGLQR (**KAP1_K779**) | 0 | **5.83x10$^{6}$** | 0 | In cellulo ubiquitination assay and K-e-GG enrichment (approach #3) |

**Table S2.** K254, K319 and K779 residues of KAP1 identified in high-throughput proteomic studies as potential ubiquitination sites.

| Site | Sequence | LTP[#] | HTP[##] |
|---|---|---|---|
| K254-ub | DAVRNQRkLLAsLVk | 0 | 26 |

| | | |
|---|---|---|
| TIF1B (human) | DAVRNQRkLLAsLVk | K254-ub |
| TIF1B iso2 (human) | DAVRNQRKLLASLVK | K172 |
| TIF1B (mouse) | DAVRNQRkLLAsLVk | K255-ub |
| TIF1B (rat) | DAVRNQRKLLASLVK | K256 |

| Site | Sequence | LTP[#] | HTP[##] |
|---|---|---|---|
| K319-ub | VLVNDAQkVTEGQQE | 0 | 112 |

| | | |
|---|---|---|
| TIF1B (human) | VLVNDAQkVTEGQQE | K319-ub |
| TIF1B iso2 (human) | VLVNDAQKVTEGQQE | K237 |
| TIF1B (mouse) | VLVNDAQkVTEGQQE | K320-ub |
| TIF1B (rat) | VLVNDAQKVTEGQQE | K321 |

| Site | Sequence | LTP[#] | HTP[##] |
|---|---|---|---|
| K779-ub | FNkLTEDkADVQsII | 0 | 49 |

| | | |
|---|---|---|
| TIF1B (human) | FNkLTEDkADVQsII | K779-ub |
| TIF1B iso2 (human) | FNKLTEDKADVQSII | K697 |
| TIF1B (mouse) | FNkLTEDkADVQsII | K779-ub |
| TIF1B (rat) | FNKLTEDKADVQSII | K780 |

[#]The number of records in which this modification site was determined using methods other than discovery MS (LTP, low-throughput).
[##]The number of records in which this modification site was assigned using high-throughput (HTP) mass spectrometry. The data are from https://www.phosphosite.org
